# High-Speed Imaging of Paw Withdrawal Reflex to Objectively Assess Pain State in Mice

**DOI:** 10.1101/263400

**Authors:** Ishmail Abdus-Saboor, Nathan T. Fried, Mark Lay, Peter Dong, Justin Burdge, Ming Lu, Minghong Ma, Xinzhong Dong, Long Ding, Wenqin Luo

## Abstract

Rodents are often used for studying chronic pain mechanisms and developing new pain therapeutics, but objectively determining the animal’s pain state is a major challenge. To improve the precision of using reflexive withdrawal behaviors for interpreting the mouse pain state, we adopted high-speed videography to capture sub-second movement features of mice upon hind paw stimulation. We identified several parameters that are significantly different between behaviors evoked by innocuous and noxious stimuli, and combined them to map the mouse pain state through statistical modeling and machine learning. To test the utility of this approach, we determined the pain state triggered by von Frey hairs (VFHs) and optogenetic activation of two nociceptor populations. Our method reliably assesses the “pain-like” probability for each mouse paw withdrawal reflex under all scenarios, highlighting the improved precision of using this high resolution behavior-centered composite methodology to determine the mouse pain state from reflexive withdrawal assays.

## Introduction

Chronic pain affects over one hundred million people in the United States, yet the mechanisms responsible for pathological pain signaling are still not fully understood. To interrogate this, reliable animal models that mimic key features of pain in humans are imperative^1^. However, it is very challenging to objectively measure pain state in rodents, as pain is a complicated and subjective experience and rodents are non-verbal.

The current assays to score pain in rodents can be broadly classified as operant pain assays, spontaneous pain detection assays, and reflexive withdrawal assays^1–4^. Operant assays typically involve animals successfully completing a task or learning to avoid or prefer an enclosed chamber that is associated with pro- or anti-nociceptive stimuli/experiences^5–8^. Since these assays require normal learning/memory processes in the animal to report its pain state, the failure of an animal to learn/remember a pro-nociceptive chamber/task may not necessarily indicate a lack of pain. Spontaneous pain detection assays, such as the grimace scale or paw licking/biting, have the advantage of mimicking the spontaneous pain that is commonly observed in the clinic^9^. Nevertheless, spontaneous measurements of pain are more difficult to quantify and not as conducive to high-throughput pre-clinical testing.

Over the past 50 years, the most widely-used measurements of pain in rodents have been reflexive withdrawal assays, in which a noxious or innocuous stimulus is applied to a region of the rodent, such as the paw or the tail, and the withdrawal frequency or latency is quantified as a readout for the animal’s pain state^2–4^. The underlying assumption for this assay is that noxious stimuli trigger “pain” sensation, whereas innocuous stimuli trigger “non-pain” sensation. Obvious advantages to reflexive assays are the ease of the procedures, the ability to test many animals in a short time period, and the similarities to human reflexes that allow for the interpretation of the results based on human experience. While these assays have led to many important discoveries in the pain field, they also have some well-recognized limitations. First, the definitions of noxious and innocuous stimuli rely on subjective human judgment, which will generate inconsistency when different research groups cannot reach a consensus on the quality of a stimulus. For example, despite the popularity of the von Frey hair (VFH) test, there is no consensus on the sensation that is triggered by VFHs in rodents^10–12^. Second, humans and rodents could have a different sensory experience to a given stimulus (i.e., a stimulus that is innocuous to humans could be noxious to rodents), so the human sensory experience may not be reliable for annotating the quality of a stimulus when it is applied to rodents. Third, there is not always a linear relationship between stimulus intensity and the experimental read-out (frequency of withdrawal reflex), as a high frequency of paw withdrawal is observed for both noxious pinprick and innocuous dynamic brush^13,14^.

One main reason that current reflexive assays in rodents utilize a combination of stimulus quality assessment and binary scoring (presence or absence of the withdrawal reflex) is because the movements occur rapidly on a millisecond scale, making it challenging to quantify and distinguish movement patterns with the unaided eye or consumer-grade cameras. Inspired by the application of high-speed videography in fly, fish, and mouse to map movement features of specific behaviors and dissect underlying neural circuits and genes^15–18^, we adopted high speed imaging (500 to 1000 frames per second (fps)) to capture movement features of the mouse paw withdrawal reflex in response to four natural mechanical stimuli (static cotton swab, dynamic brush, light pinprick, and heavy pinprick). Prior to performing high-speed behavioral analysis, we used whole animal *in vivo* calcium imaging to confirm that cotton swab and dynamic brush mainly activated intermediate and large diameter dorsal root ganglion (DRG) neurons (low-threshold mechanoreceptors for triggering “touch” sensation), whereas pinprick preferentially activated small diameter DRG neurons (high-threshold nociceptors for triggering “pain” sensation). Using these four well-defined innocuous and noxious mechanical stimuli, we characterized sub-second paw and head movement features of the withdrawal reflex in CD1 and C57 male and female mice. We identified six distinguishing movement features, which include both reflective and affective aspects of pain related behaviors, and combined them using principal component analyses to map each withdrawal reflex into “pain” vs. “non-pain” domains. We further predicted the probability of being “pain-like” for each withdrawal reflex using machine learning. To test the implication of our new approach, we applied this method and our established parameter database to study paw withdrawal in response to three VFHs. We demonstrated for the first time, to our knowledge, the sensation that is triggered by different VFHs under baseline conditions. Lastly, with this method, we revealed that acute optical activation of a broad population of nociceptors, using *TrpV1^Cre^* mediated recombination (TRPV1-ChR2 mice), led to a characteristic “painful” paw withdrawal, whereas optical activation of a more specific population of nociceptors, MRGPRD+ non-peptidergic nociceptors (MRGPRD-ChR2 mice), led to a non-painful paw withdrawal under baseline conditions. Under chronic inflammation, the same optical activation of MRGPRD+ non-peptidergic nociceptors triggered “painful” paw withdrawals, which were completely reversed to non-painful withdrawals following analgesic administration. Since TRPV1-ChR2 and MRGPRD-ChR2 mice show an indistinguishable high frequency (>70%) of paw withdrawal upon optical stimulation under all conditions, these results highlight the improved precision of our new method to annotate the mouse “pain state”. Taken together, we have developed a new method that combines high-resolution mapping of paw withdrawal movement features with statistical modeling to determine the mouse pain state. Our method should help to improve rigor and reproducibility of rodent pain research.

## Results

### *In vivo* calcium imaging to determine stimulus quality

We sought to use high-speed videography (500 to 1000 fps) to record sub-second, full-body movements of mice in response to mechanical stimuli applied to the plantar surface of the hind paw to extract behavior parameters that allow us to differentiate the mouse “pain state” (Fig. 1a). We began our analysis with four natural mechanical stimuli that are widely considered by the field as innocuous or noxious. They are static cotton swab (gently pressing a blunted, cone-shaped cotton swab against the plantar surface of the hind paw, which represents an innocuous static mechanical stimulus), dynamic brush (sweeping a soft-bristled makeup brush from the proximal to distal plantar surface, which represents an innocuous dynamic mechanical stimulus), light pinprick (gently placing a needle on the plantar surface, which represents a potentially noxious mechanical stimuli), and heavy pinprick (forcefully pushing a needle onto the plantar surface, which represents a noxious mechanical stimuli).

**Figure 1.**
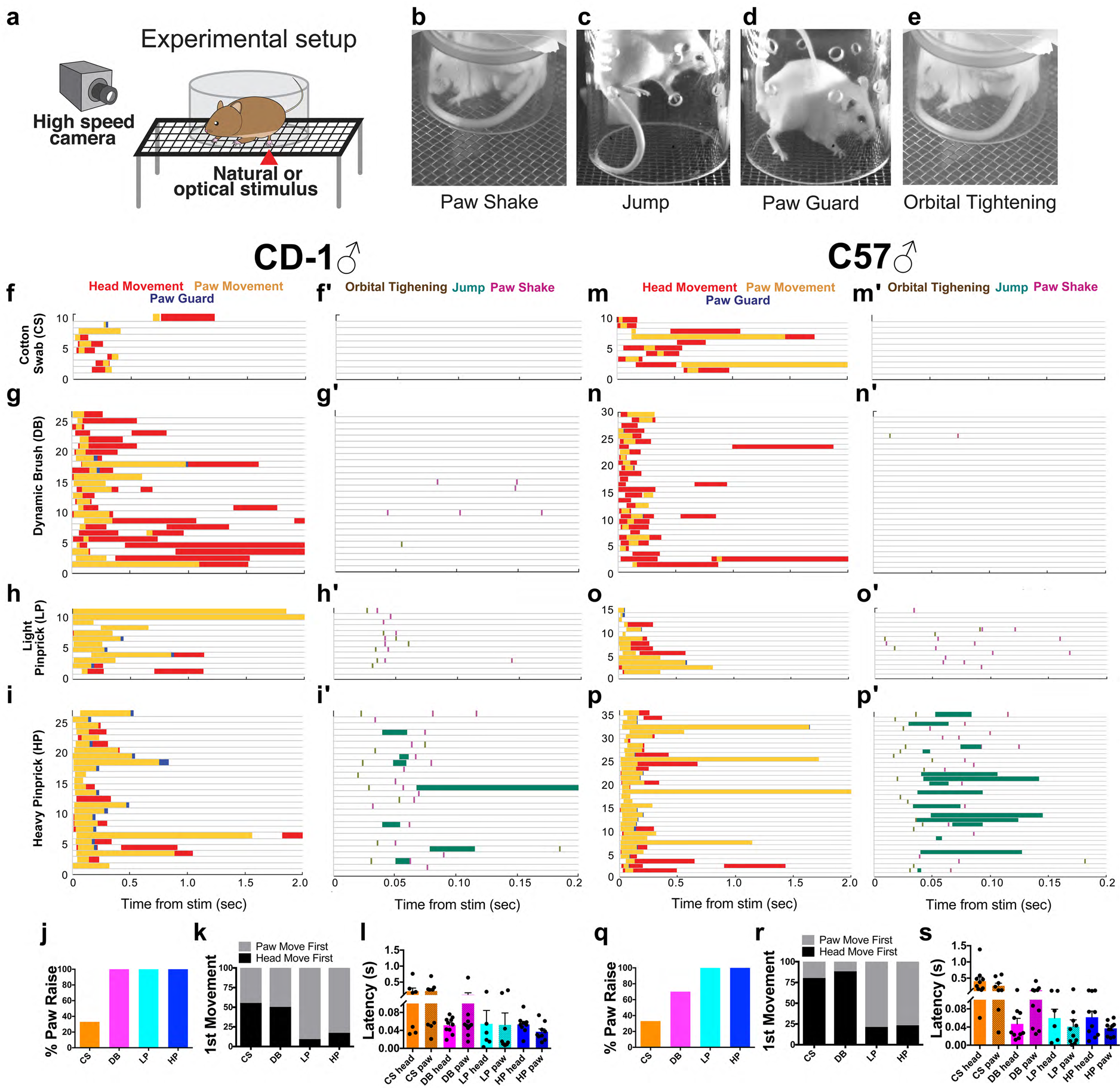
Sub-second temporal mapping of mouse behavioral features in response to paw application of natural mechanical stimuli. (a) Schematic of behavioral setup showing lateral placement of high-speed camera in relation to a contained, yet freely behaving mouse. (b-e) Representative single frame images taken from high-speed videos of CD-1 male mice following stimulus application. (f-i′ and m-p′) Responses of CD1 and C57 male mice to paw stimulation of cotton swab (CS), dynamic brush (DB), light pinprick (LP), and heavy pinprick (HP) are plotted as raster plots, showing when six behavior features (color-coded in the figure) occurred after stimulus onset within the first 2 s (f-i, m-p) or the first 200 ms (f′-i′, m′-p′). For each raster plot, the times when the behaviors occurred are shown on the X-axis, while the Y-axis and each horizontal line show a single trial/animal. Each horizontal line from the two columns of raster plots for a given strain is from the same trial. (j) Percentage of paw raise towards a given stimuli for CD1 males, n = 10. (k) First movement, whether head (black) or paw (grey), after stimulus application for CD1 males. (l) Latency of head and paw movement upon each stimulation for CD1 males. (q) Percentage of paw raise towards a given stimuli for C57 males, n = 10. (r) First movement, whether head (black) or paw (grey), after stimulus application for C57 males. (s) Latency of head and paw movement upon each stimulation for C57 males.

We first examined the sensory neuron activation patterns evoked by these four stimuli with *in vivo* calcium imaging, where we could record Ca^2+^ transients of ~1500 dorsal root ganglion (DRG) neurons per trial with the genetically encoded calcium indicator *GCAMP6* driven by the *Pirt* promoter (Supplemental Fig. 1)^19^. We applied each stimulus to the hind paw of lightly anesthetized Pirt-GCAMP6 mice in an innocuous to noxious order while recording DRG Ca^2+^ influx. Robust and rapid Ca^2+^ influx occurred within DRG neurons following the application of all four stimuli, and the number of activated neurons positively correlated with the stimulus intensity (i.e., the lowest intensity stimulus, cotton swab, activated ~5 neurons/trial and the highest intensity stimulus, heavy pinprick, activated ~15 neurons/trial (Supplemental Fig. 1a-h, m). On average, we observed Ca^2+^ transients increasing between 1 and 4 fold over baseline following the application of stimuli (Supplemental Fig. 1i-l, Supplemental Raw Data File 1). Moreover, cotton swab and dynamic brush predominantly activated intermediate (20 to 25 μm) or large (>25 μm) diameter DRG neurons, while the light and heavy pinprick stimuli predominantly activated small (< 20 μm) diameter DRG neurons (Supplemental Fig. 1n, o). These activation patterns are consistent with the notion that cotton swab and dynamic brush stimuli preferentially trigger “touch” sensation by activating large-diameter low-threshold mechanoreceptors whereas pinprick stimuli preferentially trigger “pain” sensation by activating small-diameter high-threshold nociceptors.

### High speed imaging of paw withdrawal reflex revealed distinctive movement features in response to innocuous and noxious mechanical stimuli

With confirmation about the stimulus quality, we then applied these four mechanical stimuli to the plantar surface of a randomly chosen hind paw of fully acclimated mice. To test for potential genotype- and/or sex-specific features, we examined stimulus-evoked responses in male and female CD1 and C57 wild-type mice (n = 10 for each group). All four mechanical stimuli evoked movements of the stimulated paw, the head, and the entire body, which would generally be completed within 500 ms (Supplemental Videos 1-16). We found similar patterns in both male (Fig. 1) and female mice (Supplemental Fig. 2). A typical movement sequence involved the stimulated hind paw moving away from or the head turning toward the stimulus, followed by the whole body turning. We focused on the movement features of the paw and head because they are most closely related to the stimulus onset and thus most likely reflect sensation evoked by the stimuli. The paw-associated movements usually started with the paw being raised to a maximum height. It would then be held at the apex, returned to the wire mesh, or begin a sinusoidal paw-shake (Fig. 1b). In some pinprick trials, the mouse would jump into the air with all four paws rising away from the stimulus (Fig. 1c). The mouse would then return its paw to the mesh, often in a guarding manner (only toes or heel of the paw in contact with the mesh) (Fig. 1d). The head-associated movements involved orientation/turning of the head toward the stimulus. In some instances, primarily with noxious pinprick stimuli, the mouse would display orbital tightening, which occurs in mice during pain-related grimace (Fig. 1e)^9^.

**Figure 2.**
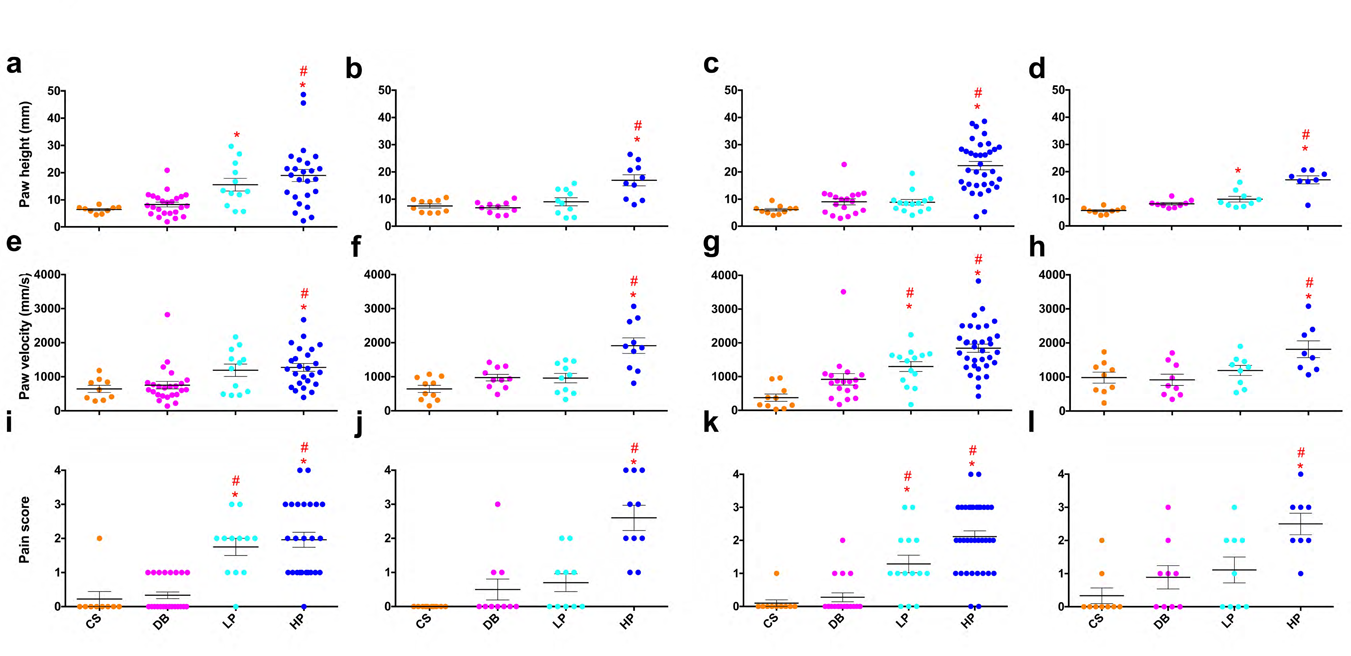
Quantification of the three most relevant behavior parameters. For all data, a single dot represents a given trial from either C57 of CD1 males. (a-d) The maximum height of the first paw raise of the stimulated paw. (e-h) The paw velocity of the first paw raise of the stimulated paw. (i-l) The pain score for a given animal to each stimulus. The pain score is a composite measurement of orbital tightening, jumping, paw shaking, and paw guarding. Statistical significance between stimuli are determined by one-way ANOVA followed by Tukey’s multiple comparison test. Red asterisks represent p-values < 0.05 when comparing CS to LP or CS to HP (LP or HP > CS), while red stars represent p-values < 0.05 when comparing DB to LP or DB to HP (LP or HP > DB). The blue asterisk indicates a statistic difference, p-values < 0.05, when comparing CS to LP (LP < CS). The error bars represents SEM, while the longest horizontal line represents the mean.

For both C57 and CD1 male mice, the likelihood of observing paw and/or head movement and the temporal order between paw/head movements depended on the stimulus type (Fig. 1). Paw movement occurred in 30-40% of trials with cotton swab for males of both genotypes, 70% in C57 and 100% in CD1 with dynamic brush, and nearly 100% for both genotypes with pinprick stimuli (Fig. 1j, q). Head movement showed the opposite trend; it occurred in 80-100% of cotton swab and dynamic brush trials but only 47-60% of light and heavy pinprick trials (Fig. 1f-p’). For both genotypes, paw movement was initiated earlier than head movement in most pinprick trials, while the order was more variable for cotton swab and dynamic brush stimuli (Fig. 1h, i, o, p, k, r). For dynamic brush, light pinprick, and heavy pinprick, the latency to the head response was ~ 50 ms for both genotypes (Fig. 1l, s). The paw response latency was also ~ 50 ms for pinprick stimuli and 100 ms for dynamic brush. The latency to paw or head movement for cotton swab was much longer, taking more than 500 ms for a response. Together, these results suggest that innocuous mechanical stimuli preferentially trigger an “exploring head turn” reflex whereas noxious mechanical stimuli are more likely to evoke a quick “avoidance paw withdrawal” reflex.

Moreover, the prevalence of certain movements, such as orbital tightening, paw shake, jumping and paw guarding (Fig. 1b-e), are closely correlated with the stimulus quality. Their incidence is rare (10%) in the cotton swab trials (Fig. 1f, f’, m, m’), occasional (15%) in the dynamic brush trials (Fig. 1g, g’, n, n’), and more frequent in the light pinprick (60%) and heavy pinprick (85%) trials (Fig. 1h-i’, 1o-p’). These behavior features were suggested to be associated with affective aspect of pain sensation^20^.

### A subset of movement parameters account for the majority of variance in the responses

To determine which movement features best distinguish between behaviors in response to innocuous and noxious stimuli, we measured a set of parameters for approximately half of trials with CD1 and C57 male mice as a pilot analysis, including: 1) the total time the paw is in movement (total paw time), 2) the total time the paw is in the air (paw air-time), 3) the total time the paw is held at the apex (paw at apex), 4) the total time the paw is in movement after reaching the apex (paw time after apex), 5) paw lift height, 6) paw lift velocity, 7) response latency (whether it be the head or paw), 8) the duration of head movement, 9) the duration of full-body movement, 10) the total behavior time, and 11) a pain score that provides a total incidence count of paw shaking, jumping, paw guarding, and orbital tightening, in a given trial. With these multi-dimensional data, we first tried to decide which of the 11 parameters could account for the majority of variance and were thus likely to be useful for distinguishing responses to innocuous vs. noxious stimuli. We found that total paw time, paw-air time, paw at apex, and paw time after apex were highly correlated (Supplementary Fig. 3a), suggesting that these four parameters were measuring the same underlying effect and contributed redundantly to the overall variance. Therefore, we focused only on paw air-time, a parameter that has been used for rodent pain behavior studies before for further analyses. We then performed an iterative exploratory factor analysis with the remaining 8 parameters (Supplementary Fig. 3b). We found that response latency, duration of head movement, duration of full-body movement, and the total behavior time either had low factor-loadings or cross-loaded onto multiple principle components, suggesting that they minimally accounted for the total variation within the system or contributed little in differentiating “pain” vs. “non-pain” responses. The last iterative exploratory factor analysis also revealed a low factor loading for paw-air time (0.205 in Supplemental Fig. 3b), suggesting that it too contributed minimally to the total variation within the system and thus is not an effective parameter for differentiating behaviors induced by each stimulus. In contrast, three parameters (paw height, paw velocity, and pain score) had high factor-loadings (0.904, 0.873, and 0.819, respectively, in the final iterative factor analysis) (Supplemental Fig. 3b) and featured an increasing trend in raw values with increasing stimulus-intensity, indicating that they likely accounted for the majority of the system’s variance and would be the most useful for differentiating between behaviors evoked by innocuous and noxious stimuli. This result was further supported in that only these three parameters had some significant differences between the behaviors evoked by innocuous versus noxious stimuli (Fig. 2) (Fig. 1 and Supplemental Fig. 2 and 4 show the other measurements). Therefore, we subsequently used these parameters, which encompass both reflexive and affective components of the pain response, to analyze withdrawal reflex behaviors from all mice (Fig. 2).

**Figure 3.**
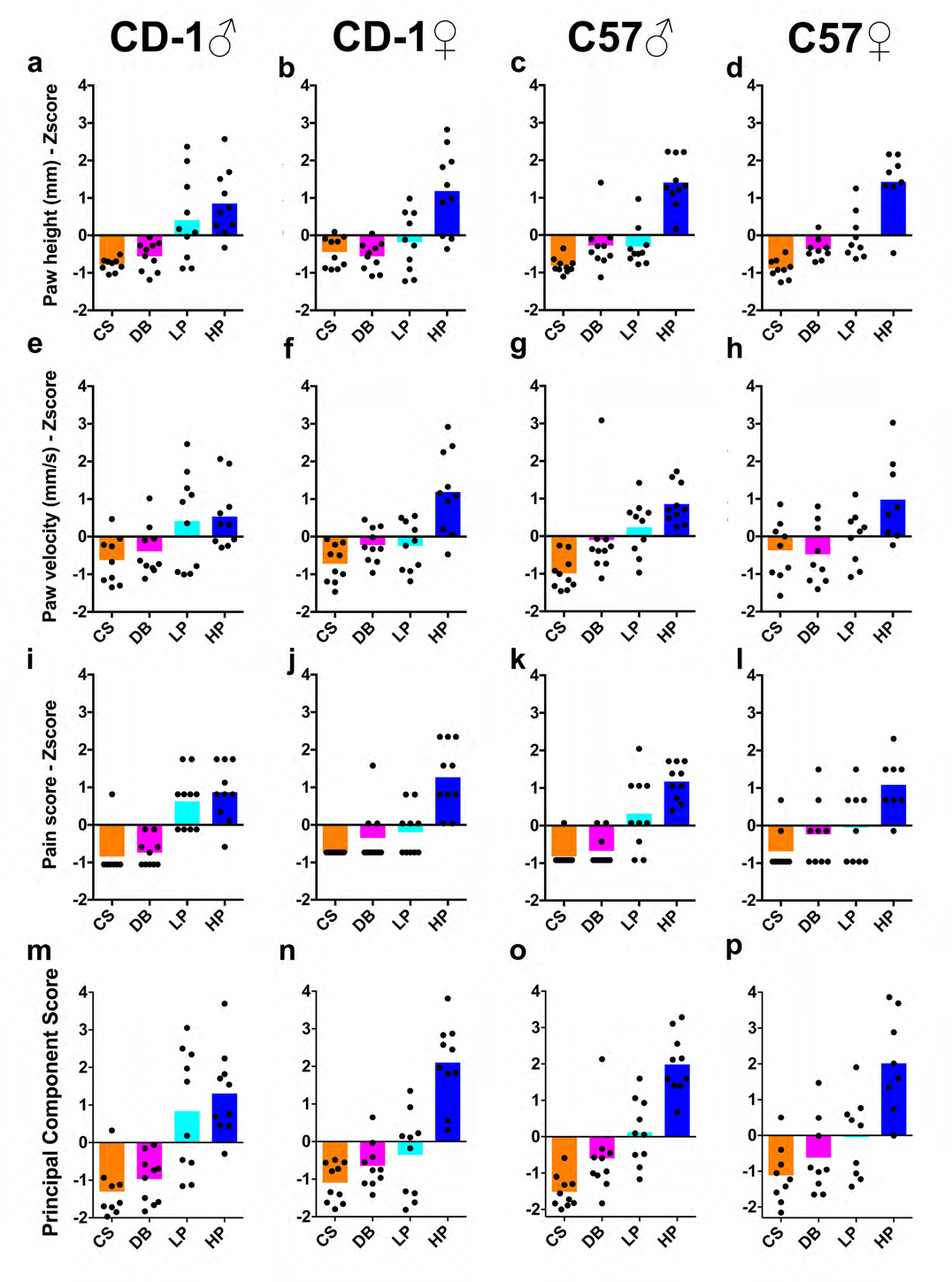
Statistical analyses to integrate four parameters into one normalized PC score. (a-l) Z-scores of individual mice are plotted relative to the combined mean from the 4 groups of sex/genotype in Figures 1, 2. Each dot represents an individual mouse. Multiple trials of the same mouse from the same stimulus were averaged first for this analysis. Plotted are Z-scores for paw height (a-d), paw velocity (e-h), and pain score (i-l). The first principal component was plotted following calculation of Z-scores for individual measures and obtaining eigenvalues (m-p) (see Methods and Supplementary Table 1).

**Figure 4.**
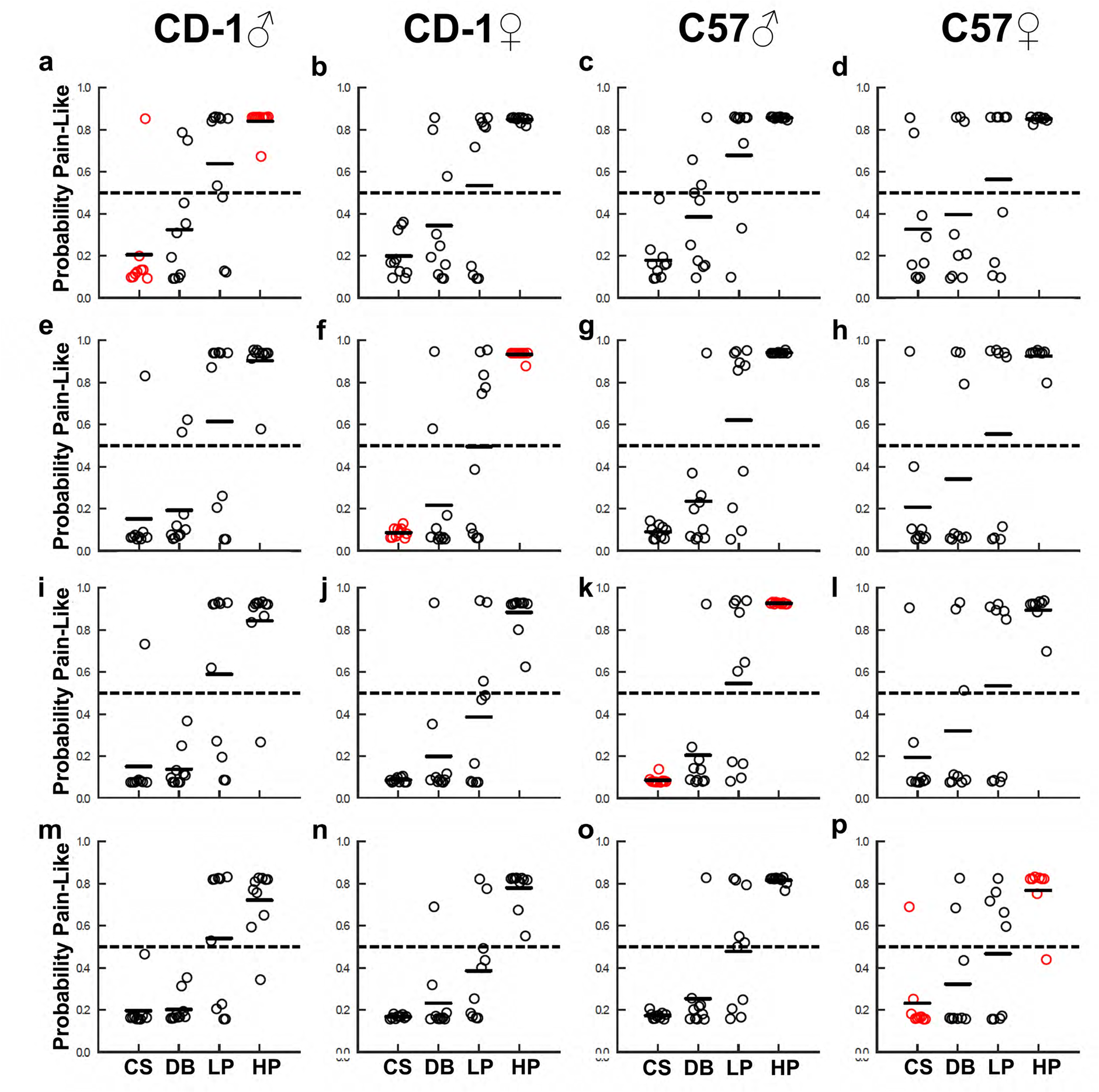
Machine learning predicts “pain-like” probability for each paw withdrawal reflex. Trained support vector machine (SVM) analyzed each behavior trial and output its probability of being pain-like. (a - d) Predictions made following training with CS and HP trials from CD1 males. (e - h) Predictions made following training with CS and HP trials from CD1 females. (i - l) Predictions made following training with CS and HP trials from C57 males. (m - p) Predictions made following training with CS and HP trials from C57 females. Red circles denote trials used for the training of the SVM. Each dot represents the averaged response of an individual mouse.

### A combination of distinguishing movement parameters can indicate mouse pain state

Although each of the three parameters provide information that helps distinguish movements induced by noxious vs. innocuous stimuli, they are expressed in different dimensions with regard to absolute values and units. In addition, it is unclear what the exact “pain vs. non-pain” threshold is for each parameter. To take advantage of the entire set of information, we sought to combine these three different parameters into a one-dimensional score using a principal component analysis (PCA): first converting the raw data to normalized Z scores within each dataset (Fig. 3a-l), and then applying a PCA on converted Z scores to determine the relative contribution for each parameter (as reflected by eigenvalues, see Supplemental Table 1). The first principal component score (PC1 score) of the three-dimensional dataset was computed as a weighted total value. To account for potential genotype/sex differences, the PCA was performed separately for the four genotype-sex combinations, generating four separate sets of eigenvalues to calculate PC scores.

With this transformation, we were able to plot the PC scores for reflex behaviors in response to each stimulus within males and females of both genotypes (Fig. 3m-p). We found that: 1) different from the withdrawal frequency, mean PC scores were positively correlated with increasing stimulus intensity (i.e., PC scores for cotton swab < dynamic brush < pinprick) in males and females of both genotypes, such that higher intensity noxious mechanical stimuli (heavy pinprick (blue)) which triggered “pain” sensation, results in mostly positive PC scores while lower intensity innocuous mechanical stimuli, such as cotton swab (orange) and dynamic brush (magenta) which trigger “non-pain” sensation, results in mostly negative PC scores; 2) mean PC scores for light pinprick trials (cyan) were the most variable across genotype and sex (i.e., PCs scores were positive for most CD1 male trials but negative for most CD1 female trials, suggesting that light pinprick may trigger a different sensation in male and female mice); and 3) for a given stimulus-type in a given strain/sex combination, there was considerable variation in PC scores among different mice, which may be caused by variations of the internal state of each animal during testing (i.e., alert, resting, etc.) or the slight stimulus variability from trial to trial. Taken together, our PC-based analysis suggests that the PC score of each mouse can be used to map its individual “pain state” and intensity, with a score of “0” serving as the potential threshold to separate pain versus non-pain domains.

### Machine learning classifies withdrawal behaviors as a probability of being pain-like

To further classify mouse pain state based on their reflexive behaviors, we used a machine-learning approach to predict the probability of each trial being pain-like. Specifically, we used the PC scores of cotton swab and heavy pinprick trials from one group of mice to train a support vector machine (SVM) (Supplemental Table 2). Cotton swab and heavy pinprick trials were chosen because their triggered behaviors can be defined as “non-pain” or “pain” with high confidence and the corresponding PC scores showed the most consistent patterns across genotype/gender. The trained SVM was then used to predict the probability of being “pain-like” for all trials.

We first determined the predicted pain-like probability for withdrawal reflex triggered by dynamic brush or light pinprick within the same genotype/sex group (Fig. 4, red circles indicate the training data). To do this, the SVM was trained with cotton swab and heavy pinprick data from CD1 males (Fig. 4a), CD1 females (Fig. 4f), C57 males (Fig. 4k), or C57 females (Fig. 4p). The pain-like probabilities for behaviors triggered by dynamic brush ranged from 0.20 to 0.33 and for light pinprick ranged from 0.47 to 0.65 (Fig. 4a, f, k, and p). Thus, these results suggest that dynamic brush had a low probability of evoking pain-like sensation (< 0.33) in each sex of both genotypes, despite the ~100% responsive rate. Notably, the SVM predictions revealed that only responses of CD1 males to light pinprick had a high probability of being pain-like (0.65). In all other groups, the probability was no greater than 0.55. Thus, similar to PC scores, these results suggest that mice with different genetic backgrounds or sex may sense light pinprick as noxious or innocuous.

Given the known effect of genetic background and sex on pain sensation, we next asked whether a SVM trained with cotton swab and heavy pinprick from one sex and genotype could reliably classify similar trials from the other sexes/genotypes. Under these training conditions, we found a consistently high pain-like probability for heavy pinprick trials (range of 0.69 to 0.96) and a low pain-like probability for cotton swab trials (range of 0.08 to 0.28). Dynamic brush was also consistently predicted to have a low probability of being pain-like (range of 0.14 to 0.39) while light pinprick was consistently predicted to have a boundary probability of being pain-like (range of 0.39 to 0.68). Notably, these predictions for dynamic brush and light pinprick, when trained with a different sex or genotype, are more variable than the predictions made when training with the same sex and genotype. Thus, SVM trained with cotton swab and heavy pinprick data from one sex/genotype group could be used to reliably classify responses to the same stimuli from another group. Classification of responses to other stimuli, such as dynamic brush or light pinprick, however, work best with training data sets from matched sex/genotype group.

### High-speed imaging analysis of paw withdrawal reflex triggered by von Frey hairs

We next sought to validate the usefulness of our approach by analyzing the paw withdrawal reflex of CD1 male mice in response to three VFHs (0.6 g, 1.4 g, and 4.0 g). These filaments are often used to measure mechanical threshold or mechanical pain responses in mice^21^. Although each VFH delivers a well-defined amount of mechanical force, whether it triggers an innocuous or noxious responses for a mouse under a given condition is uncertain.

CD1 male mice responded to 50% of 0.6 g VFH trials, 90% of 1.4 g VFH trials, and 100% of 4.0 g VFH trials (Fig. 5a) (Supplemental Videos 17-19), similar to what is reported in the literature ^14,21^. Paw height was significantly greater for 4.0 g VFH compared to 1.4 and 4 g VFH (p = 0.004 and 0.027, respectively) (Fig. 5b). Likewise, paw velocity was also significantly greater for 4.0 g VFH compared to 1.4 and 4 g VFH (p < 0.0001 and 0.012, respectively) (Fig. 5c). Conversely, no statistical difference (p > 0.215) in pain score was found between any of the filaments (Fig. 5d). The PC score of each response to a given VFH was calculated using the Z scores of the three parameters and the previously obtained eigenvector values from CD1 male data (Supplemental Table 1). On average, PC scores were positive (0.246) for 4.0 g VFH and negative for 0.6 g (-0.957) and 1.4 g (-0.498) (Fig. 5e). Additionally, the SVM generated from and trained with cotton swab and heavy pinprick data from CD1 males predicted a high pain-like probability for the 4.0 g VFH (0.80), a low pain-like probability for the 0.6 g VFH (0.33), and a boundary pain-like probability for the 1.4 g VFH (0.51) (Fig. 5F). Taken together, the analysis using our new method reveals that the 4.0 g VFH filament likely evokes a pain-like withdrawal reflex, the 0.6 g VFH likely evokes a non-pain withdrawal reflex, while 1.4 g may be near the mechanical threshold separating pain from non-pain responses. As far as we know, this is the first clear “sensory quality interpretation” for withdrawal reflex triggered by different VFHs.

**Figure 5.**
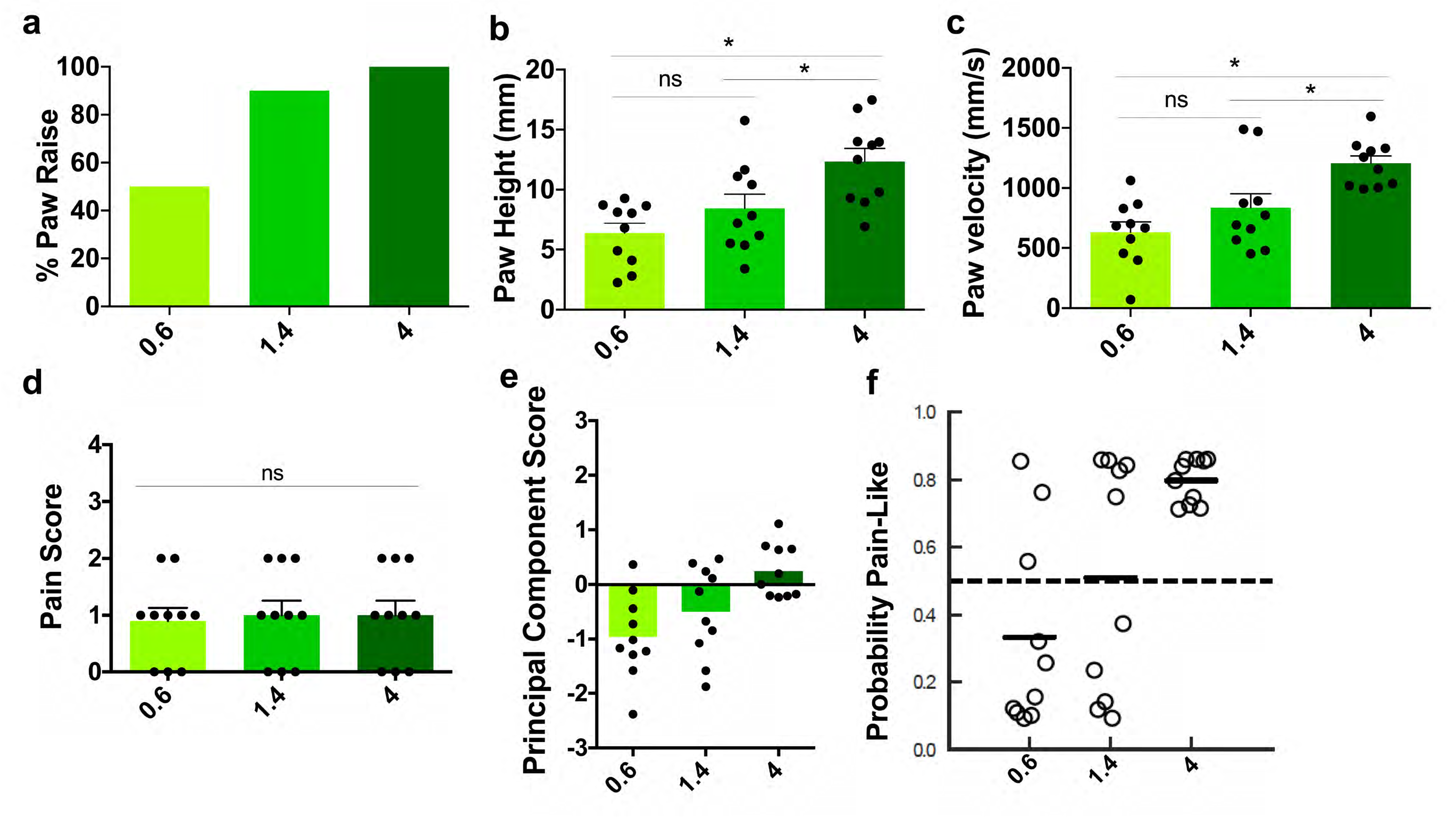
Analysis of mouse paw withdrawal reflex in response to von Frey hairs. (a) Responsive rate for each VFH filament. (b) Paw height, (c) paw velocity, (d) and pain score were quantified for each VFH filament. (e) Principal component score plot for each VFH filament. (f) SVM predications for each VFH filament. SVM was trained with CS and HP data of CD1 males. N = 10 CD1 male mice.

### High-speed imaging analysis of paw withdrawal reflex triggered by peripheral optogenetic activation of different primary afferent populations

Optogenetics is a powerful gain-of-function approach to study primary somatosensory afferents ^18,22–25^. Briefly, channelrhodopsin (ChR2) is expressed in different DRG neuronal populations and application of transdermal light is used to activate ChR2+ afferents in the skin. Interestingly, optogenetic activation of different populations of DRG neurons reported in the literature thus far all triggered paw withdrawal reflex, raising the question of how to interpret the meaning of the paw withdrawal when using peripheral optogenetics.

Here we tested whether the high-speed imaging and statistical analysis method we established using wild type mice and natural mechanical stimuli could be applied to the analysis of light-induced withdrawal behaviors of transgenic mice. For this purpose, we generated two mouse lines. For the first line (TRPV1-ChR2 mice), we crossed *TrpV1^Cre^* ^26^ to the Ai32 Cre-dependent ChR2 line^27^ to express ChR2 in the majority of nociceptors (91% of CRGP+ and 95% of IB4+ nociceptors) and a few large diameter DRG neurons (13.4% of NFH+ mechanoreceptors (Supplemental Fig. 5). For the second line (MRGPRD-ChR2), we crossed an inducible Cre mouse line generated in our lab *Mrgprd^CreERT2^* ^28^, to Ai32 in which ChR2 is specifically expressed in non-peptidergic MRGPRD+ C-nociceptors (Supplementary Fig. 6).

**Figure 6.**
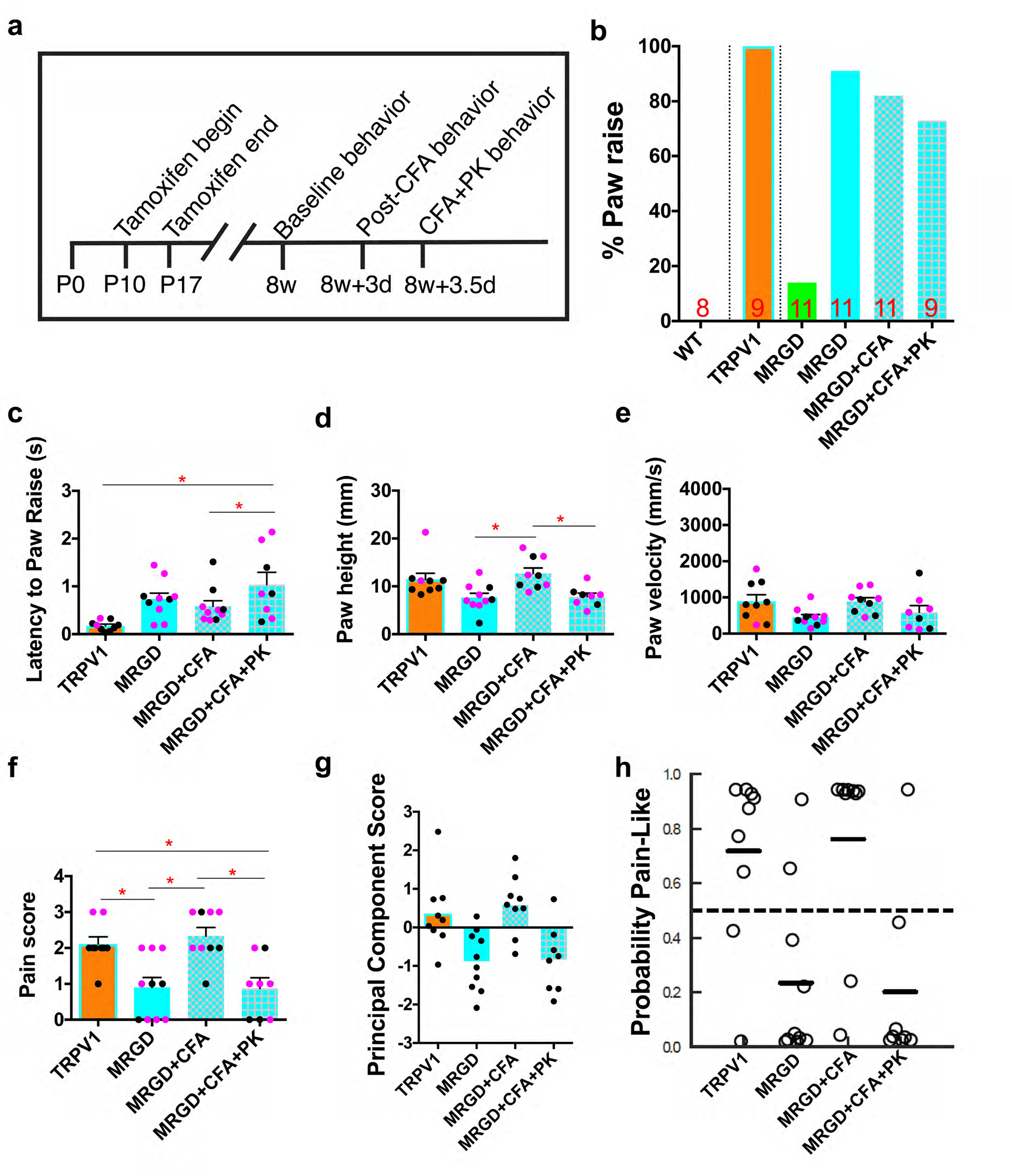
Optogenetic activation of MRGPRD^+^ neurons leads to “pain-like” or “non-pain” paw withdrawal reflex depending on the animal state. (a) Diagram showing the treatment paradigm and experimental design for paw reflexive behavior assays with MRGPRD-ChR2 mice. (b) Percentage of animals displaying a paw raise. WT indicates *ChR2^f/f^* littermate control, orange bars indicates TRPV1-ChR2 mice at baseline, blue bars indicates MRGPRD-ChR2 mice at baseline before CFA injection, 3 days post CFA, and painkiller injection at 3.5 days after CFA. (c) Latency between first blue light stimulation and paw raise. (d-f) quantification for paw height (d), paw velocity (e), pain score (f). (g) PC scores of TRPV1-ChR2 and MRGPRD-ChR2 mice at baseline, after CFA, and CFA + painkillers using eigenvectors derived from wild type mice of both sexes and genotypes. (h) SVM pain-probability graphs using all wild type mice of both sexes and genotypes as training datasets, to predict the probability of a pain response for TRPV1-ChR2 and MRGPRD-ChR2 optogenetic responses in baseline, after CFA, and CFA + painkillers.

Hind paw stimulation with blue laser light (10 mW) of ChR2-only littermate control mice did not cause paw withdrawal, as we have previously reported^28^ (Fig. 6a, b) (Supplemental Video 20), suggesting that the blue laser stimulation itself does not cause non-specific sensation. Blue laser light hind paw stimulation of TRPV1-ChR2 mice induced a paw withdrawal in 100% of mice with a response latency of ~150 ms (Fig. 6b, c) (Supplemental Video 21). Paw height, velocity, and pain score were quantified as previously established (Fig. 6d-f). Since these transgenic mice are on a mixed C57 and CD1 genetic background and contain both males and females, we calculated PC scores using eigenvalues derived from a combined dataset of C57 and CD1 wild type male and female mice (Fig. 6g, Raw Data File 2, and Supplemental Table 1) and predicted the “pain-like” probability using an SVM trained with CS and HP from male and female CD1 and C57 mice (Fig. 6h). We also did analysis in a sex-specific manner (Supplemental Fig. 7, Supplemental Table 1 & 2). Overall, our results revealed that optogenetic activation of TRPV1-ChR2+ afferents triggered paw withdrawal reflexes with positive PC scores (Fig. 6g) and a high probability of being “pain-like” (Fig. 6h).

Next, we examined behavioral responses induced by blue laser light stimulation of MRGPRD-ChR2 mice. Similar to TRPV1-ChR2, we found that 91% of MRGPRD-ChR2 mice displayed a paw withdrawal under blue laser light stimulation (Fig. 6b) (Supplemental Video 22). Most (6 out of 7) MRGPRD-ChR2 mice had no response under green laser light stimulation (Fig. 6b). The blue light-triggered paw response in MRGPRD-ChR2 mice had a latency of ~700 ms (Fig. 6c), which is ~15 times slower than natural mechanical stimuli and 4-5 times slower than blue-light-induced responses in TRPV1-ChR2 mice. Paw height, velocity, and pain score were quantified and PC scores were calculated in a similar manner as TRPV1-ChR2 mice (Fig. 6d, Supplemental Table 1, Raw Data File 2). Interestingly, PC scores of the MRGPRD-ChR2 mouse paw withdrawal reflex in response to blue laser were, on average, negative (-0.873) (Fig. 6g). The SVM also predicted a low probability that these responses were pain-like (Fig. 6h). The results were similar when the PCA1 was generated or the SVM trained in a sex specific manner (Supplemental Fig. 7). Taken together, these results show that acute activation of MRGPRD^+^ DRG neurons under baseline conditions robustly triggers paw withdrawal responses, but not “pain-like” behaviors/sensations.

To examine whether activation of MRGPRD+ neurons can trigger “pain” sensation under other conditions, we induced chronic inflammation in one hind paw of control and MRGPRD-ChR2 mice by injecting hind paws with complete Freund’s adjuvant (CFA) as previously described^29^ (Fig. 6a). We found that, although blue laser triggered a similarly high rate of responses, paw velocity, paw height, and pain score differed significantly between baseline and CFA conditions (Fig. 6c-f). The PC scores (0.578) and SVM predictions both suggest that activation of MRGPRD^+^ neurons under chronic inflammation evoked a “pain-like” withdrawal reflex (Fig. 6g,h) (Supplemental Video 23). Subsequent administration of meloxicam (2 mg/kg) and buprenorphine (0.5 mg/kg), commonly used anti-inflammatory and opioid-like analgesics^30^, inverted PC scores (-0.842) and SVM predictions to the non-pain domain, without affecting response frequency (Fig. 6b, g, and h) (Supplemental Video 24). Together, these results suggest that optogenetic activation of non-peptidergic nociceptors induces pain under inflammatory conditions. Our findings not only highlight the interesting physiology of this population of DRG neurons, but demonstrate the utility of our method to determine the pain state of mice with light-induced somatosensory behaviors.

## Discussion

We present here a novel method combining high-speed videography and statistical modeling to objectively interpret sensations associated with the mouse paw withdrawal reflex. Compared to the traditional measurements (scoring of withdrawal versus no withdrawal or quantification of withdraw latency), our approach quantifies six different behavior parameters on a sub-second scale and combines them to assess the mouse pain state. With machine learning, we are able to further determine the probability that a given paw withdrawal behavior is “pain-like” on a trial-by-trial basis. In short, this new approach would greatly improve our ability to use the rodent paw withdrawal reflex as a behavioral readout for “pain” sensation to study underlying cell and molecular circuit mechanisms or screening for new therapeutics that modulate pain.

### Development of a new method to quantify mouse pain state with improved objectivity and precision

A major concern in the pain research field is that only 11% of pain therapeutics entering Phase 1 clinical trials ever become approved by the US Food and Drug Administration^31^. Although many factors likely contribute to this low success rate, one concern is whether pain was accurately assessed in preclinical animal models, which heavily rely on reflexive behavioral assays^32^. The withdrawal reflex rate is interpreted as an indication of a “painful” or “non-painful” sensation based on the experimenter’s own subjective experience/judgement of the stimulus quality and animal state, which inevitably introduces ambiguity and potential bias that may impact the translation of preclinical findings. To address this issue, we developed a “behavior-centered” method, which allows the behavior itself to indicate the animal’s experience.

We first stimulated wild type mice with four commonly used mechanical stimuli, whose qualities were verified using *in vivo* calcium imaging (Supplementary Fig. 1). With a combination of high-speed imaging, statistical modeling, and machine learning, we characterized the detailed movement features of the paw withdrawal reflex in response to innocuous or noxious mechanical stimuli to distinguish between “painful” from “non-painful” paw lifts (Fig. 1 and Supplemental Fig. 1). The non-pain lift also features the orientation of the head toward the stimulus prior to the movement of the paw, as opposed to the pain-like withdrawal reflex that most often features movement of the paw first. This may reflect the engagement of spinal cord circuits that remove the paw from potential danger before the engagement of supraspinal circuits that would alert and direct the animal’s attention toward the stimulus^33^.

It is interesting that the PC scores, a weighted total combining relative normalized values from all six movement features together through statistical modeling, are distributed along a stimulus/sensation spectrum, with positive scores correlating to “pain-like withdrawal” and negative scores correlating to “non-pain lift” (Fig. 2,3). Our results suggest that these PC scores may be used as a scale to determine the mouse pain state, similar to the pain rating systems used for human pain assessment^34^. Further, we found that the SVM predictions, a machine learning method, can be used to predict the probability a particular withdrawal is “pain-like” after training with cotton swab (“non-pain”) and heavy pinprick (“pain”) trials (Fig. 4). This predication is accurate within the same stimulus categories, regardless of genetic backgrounds and sexes, demonstrating that this method may be useful cross-strain and cross-sex. It is also notable that these PC scores and SVM probabilities display individual variability even among the same stimulus, genetic background, and sex. This may be due to the fact that the internal state at the time of testing and the delivery of a given somatosensory stimulus would vary from trial to trial, despite identical genetic background and sex of mice. Thus, our approach enables the interpretation of sensation independently of presumptions about the stimulus quality and could be used to determine the pain status at the individual level.

### High-speed videography increases the resolution of movement features associated with distinct somatosensory stimuli

Our work adds to a very short list of papers using high-speed videography in rodents to map the movements following stimulus application to the paw^18,35,36^. Mitchell et al. used 500 fps recordings of rat hind paw withdrawals from an infrared laser, and identified paw shaking, orientation of the head toward the stimulus, and paw guarding with their analysis^35^. Further, they reported the presence or absence of these behaviors in an ordinal rating system for the assessment of pain intensity. Browne et al. used 1000 fps recordings to determine how mouse body position impacts the response to single-unit optogenetic activation of nociceptors, and focused on the latency to vibrissae, body, and paw movement^18^. Similarly, Blivis et al. used 500 fps to measure the timing of body and paw movements of rats induced by a noxious stimulus, uncovering a gating mechanism for these movements when the animal was on only its hind paws^36^. Our findings push beyond these elegant studies by not only identifying the presence/absence of “pain” associated sub-second behavior features, but by developing new statistical methods to integrate multiple relevant behavior parameters that allow us to quantitatively access mouse pain status (Figs. 3 & 4). Compared to the traditional method, the employment of high-speed imaging and this integrated scoring system greatly improves precision and confidence in annotating the mouse “pain” state.

### Proof of Concept Case 1: Von Frey hairs can induce “non-pain” or “pain-like” paw withdrawals

The VFH test is one of the most widely-used somatosensory assays^2^. However, at present there is little consensus about the sensory quality of each VFH in model organisms. Here we used our method in CD1 male mice to understand the sensory experience induced by three different VFHs (Fig. 5). We found that the 4.0 g VFH induced a “pain-like” paw withdrawal, and thus is likely to be a noxious stimulus for these mice. On the other hand, 0.6 g and 1.4 g induced withdrawal behaviors more similar to those induced by cotton swab and dynamic brush, suggesting that they are likely innocuous mechanical stimuli. Notably, mice showed similar pain associated features (i.e., pain score) in response to all three VFHs (Fig. 5d, Supplemental Fig. 8). Thus, without using high-speed imaging and our composite principal component analysis, it would be challenging to distinguish the “quality” of these responses. Additionally, although 0.6 g is often considered as the 50% “pain” withdrawal threshold^37–40^, our analysis suggests that it is well under the pain threshold (PC score of “0” and “pain-like” probability of “50%”). Instead, PC scores from 1.4 g trials are very close to the “0” threshold and have an ~50% probability of being “pain-like”, suggesting that the 1.4 g VFH is close to the threshold that separates touch and mechanical pain in mice. Our results are interesting in light of recent genetic studies ablating the mechanosensitive ion channel PIEZO2, which is critical for touch sensation^41^. When *Piezo2* is deleted from all DRG neurons and Merkel cells, a deficit in VFH responsiveness is only observed at 3.0 g and below^42^. When *Piezo2* is deleted in Merkel cells only, a deficit in VFH responsiveness is observed at 1.5 g and below^43^. Together, these ablation studies place the threshold that separates “pain” from “non-pain” at approximately 1.5-3.0 g, which is remarkably similar to what is indicated by our new method where only 4.0 g is classified as “pain-like”. To the best of our knowledge, our behavioral platform is the first to objectively demonstrate what sensation each VFH actually triggers in mice.

### Proof of Concept Case 2: Analysis of paw withdrawal reflex evoked by peripheral optogenetic approach

Non-peptidergic MRGPRD^+^ nociceptors are a molecularly and anatomically unique class of small diameter primary somatosensory neurons^44^. They are polymodal high-threshold C fibers responsive to mechanical, chemical, and thermal stimuli based on physiological recordings^45–47^. In addition, genetic ablation studies suggest that these primary afferents are tuned for detecting noxious mechanical stimuli^48^. Paradoxically, however, when non-peptidergic nociceptors were acutely activated by either chemogenetics or optogenetics using place preference assays, no place aversion was observed^49,50^. These gain-of-function studies raise the question of whether acute activation of these neurons *in vivo* is sufficient to trigger pain sensation. Here we analyzed paw withdrawal reflex upon acute peripheral optogenetic activation of ChR2^+^ MRGPRD^+^ neurons using our new method. Although acute activation of this neuronal population leads to almost 100% of paw withdrawal at baseline conditions, our PCA and SVM analyses indicates that these withdrawals fall into the domain of being “non-painful” (Fig. 6). This is in great contrast to the light evoked “pain-like” paw withdrawal reflex of TRPV1-ChR2 mice, in which ChR2 is expressed in a broader population of nociceptors. The results from TRPV1-ChR2 mice indicates that “pain” sensation can be triggered by peripherally stimulating transgenic mice expressing ChR2 with light, while the results from MRGPRD-ChR2 mice, which contains two copies of ChR2 conditional alleles and is even stimulated with a higher laser power, likely reflects the true biological functions of these neurons. Our result is in agreement with previous operant assays (chamber preference studies) ^49,50^, which suggest that, under baseline conditions, acute activation of MRGPRD+ non-peptidergic nociceptors is not sufficient to evoke “pain” sensation. This is also consistent with human self-report of “tingling” but not “pain” sensation after taking beta-alanine, a chemical that activates MRGPRD^51^. Interestingly, the VFH mechanical forces used in these previous loss-of-function study were 1.2 g and below^48^. Since our new data indicates that 1.4 g is close to the mouse mechanical pain threshold under baseline conditions, results using 1.2 g may indicate a change in the sense of touch but not necessarily mechanical pain. Collectively, these studies (both loss-of-function and gain-of-function) suggest that while non-peptidergic neurons may normally play a role in mechanical sensation, acute activation of only this population is insufficient to trigger “pain” sensation at the baseline condition.

Can MRGPRD^+^ nociceptors mediate “pain” sensation under any other conditions? Interestingly, when we used CFA to induce chronic inflammation in the mouse paw (a chronic pain model), we did observe a “pain-like” response to optogenetic activation of these afferents, as indicated by the PCA and SVM (Fig. 6). This was completely reversed by analgesic treatment (Fig. 6). Our result is congruent with the loss-of-function data, where mice without non-peptidergic nociceptors displayed much lower mechanical allodynia after chronic pain induction^48^.

Moreover, we noticed that for both TRPV1-ChR2 and CFA-injected MRGPRD-ChR2 mice, the high probabilities of light inducing pain was driven mainly by the contribution of the pain score (i.e., orbital tightening, paw shaking, jumping, and paw guarding) but not paw height or velocity (Fig. 6 & Supplemental Fig. 8). This is in contrast to the 4.0 g VFH where the high probabilities of being pain-like are driven by the contribution of paw height and velocity (Fig. 2, 3; Supplemental Fig. 8). Though the neuronal mechanisms underlying these differences are not fully understood yet, our results highlight the complexity of "pain expression phenotypes" in animals and the strength of including parameters indicating both reflective (paw height and velocity) and affective (pain score) aspects of pain for analysis^20^.

In summary, these experiments demonstrate the improved precision of our new quantitative approach with high-speed videography, which would be a vital tool in deciphering the meaning behind paw withdrawal behaviors. As more labs use peripheral optogenetic approaches to study the somatosensory system and neural circuits underlying pain sensation, our results also bring caution against presumptions for interpreting light-induced behaviors that are mainly based upon the neuronal population that expresses ChR2.

## Future Directions

In conclusion, we present here a new method combining high-speed imaging and statistical modeling to analyze the paw withdrawal reflex for interpreting the pain state of mice. One drawback of our current methodology is the reliance on manual annotation of the high-speed videos, which is time-intensive and potentially error prone. In future versions of this platform, we aim to automate the quantification and annotation process of the high-speed videos to increase analysis speed and reduce potential human error. In addition, the prediction precision will be increased when more datasets (different genetic background and sex) are used for SVM training, which is particularly important for analyzing transgenic mice that are usually in a mixed genetic background (Fig. 6). Our study here generated a database of pain-associated behavior parameters for ten CD1 and C57 male and female mice (Supplemental Raw Data file 2), which represents only an initial dataset for this method. Future studies with more animals and additional genetic background/sex would improve the robustness of the SVM predictions. Therefore, one future direction is to make this platform an open access website, where any interested lab could deposit their behavior videos and perform the analysis online. As more behavior data is collected and analyzed, the more precise and powerful this SVM prediction will become, and the more useful this will be for the whole rodent pain research field.

## Materials and Methods

### Mouse strains

Mice for behavior testing were maintained in a conventional animal facility in the John Morgan building at the University of Pennsylvania. Mice for *in vivo* calcium imaging were maintained in a barrier animal facility at the Johns Hopkins School of Medicine. All procedures were conducted according to animal protocols approved by the university Institutional Animal Care and Use Committee (IACUC) and in accordance with National Institutes of Health (NIH) guidelines. CD1 male and female mice were purchased from Charles River Laboratories and C57BL/6 male and female mice were purchased from Jackson Laboratories. MRGPRD-ChR2 mice, *Mrgprd^CreERT2^;ChR2^f/f^* (Ai32 homozygous), were generated in our lab as previously described^28^. MRGPRD-ChR2 mice were treated with tamoxifen between P10-P17 to induce robust ChR2 expression in MRGPRD^+^ neurons. TRPV1-ChR2 mice, *TrpV1^Cre^;ChR2^f/+^* (Ai32 heterozygous), were generated by crossing *TrpV1^Cre^* and Ai32 together. Pirt-GCAMP6 mice were generated by crossing *Pirt^Cre^* mice to *Rosa-GCAMP6* mice, as previously described^19^. *TrpV1^Cre^* (stock no. 017769)^26^ and *Rosa-ChR2-eYFP* (Ai32) (stock no. 012569)^27^ mice were purchased from the Jackson Laboratories.

### Whole animal L4 DRG neuron calcium imaging combined with hind paw stimulation

The L4 DRG of Pirt-GCAMP6 mice was surgically exposed and imaged, with subsequent data analysis performed using ImageJ (NIH) as previously described^19^. Briefly, for all imaging experiments, mice at 2 months or older were anesthetized by i.p. injection of chloral hydrate (560 mg/kg). After deep anesthesia was reached, the animal’s back was shaved and aseptically prepared. Ophthalmic ointment (Lacrilube; Allergen Pharmaceuticals) was applied to the eyes to prevent drying. During the surgery, mice were kept on a heating pad (DC temperature controller, FHC) to maintain body temperature at 37 ± 0.5 degrees Celsius as monitored by a rectal probe. Dorsal laminectomy in DRG was performed usually at spinal level L6 to S1 below the lumbar enlargement (occasionally at lower than S1) but without removing the dura. A 2-cm-long midline incision was made around the lower part of the lumbar enlargement area; next, the paravertebral muscles were dissected away to expose the lower lumbar part which surrounds (L3–L5) vertebra bones. The L4 DRG transverse processes were exposed and cleaned. Using small rongeurs, we removed the surface aspect of the L4 DRG transverse process near the vertebra to expose the underlying DRG without damaging the DRG and spinal cord. Bleeding from the bone was stopped using styptic cotton. After surgery, mice were laid down in the abdomen-down position on a custom-designed microscope stage. The spinal column was stabilized using custom-designed clamps to minimize movements caused by breathing and heart beats. *In vivo* imaging of whole L4 DRG in live mice could be performed for 1–6 hr immediately after the surgery.

The four stimuli (cotton swab, dynamic brush, light pinprick, and heavy pinprick) were applied to the freely hanging hind paw as described in the following section. The microscope stage was fixed under a laser-scanning confocal microscope (Leica LSI microscope system), which was equipped with macrobased, large-objective, and fast EM-CCD camera. Live images were acquired at typically eight to ten frames with 600 Hz in frame-scan mode per 6–7 s, at depths below the dura ranging from 0 to 70 mm, using a 5X 0.5 NA macro dry objective at typically 512 X 512 pixel resolution with solid diode lasers (Leica) tuned at 488 and at 532 nm wavelength and emission at 500–550 nm for green fluorescence. DRG neurons were at the focal plane, and imaging was monitored during the activation of DRG neuron cell bodies by peripheral stimuli. The imaging parameters were chosen to allow repeated imaging of the same cell over many stimuli without causing damage to the imaged cells or to surrounding tissue. Raw image stacks (512 X 512 to 1024 X 1024 pixels in the x–y plane and 20–30 mm voxel depth; typically 10 optical sections) were imported into ImageJ (NIH) for analysis. A neuron displaying Ca^2+^ ΔF/F_0_ higher than 20% is considered as a positively responsive neuron.

### High-Speed Videography

To capture mouse kinematic movement features at high temporal resolution, we recorded behaviors at 500 or 1000 frames per second (fps) with a high-speed camera (FastCAM UX100 800K-M-4GB - Monochrome 800K with 4GB memory) and attached lens (NikonZoom Wide Angle Telephoto 24-85mm f2.8). With a tripod with geared head for Photron UX100, the camera was placed at a ~45° angle at ~1-2 feet away from the Plexiglas holding chambers where mice performed behaviors. The camera was maximally activated with far-red shifted 10 mW LED light that mice cannot detect and thus would not disturb their behaviors. All data were collected and annotated on a Dell laptop computer with FastCAM NI DAQ software that is designed to synchronize Photron slow motion cameras with the M series integrated BNC Data Acquisition (DAQ) units from National Instruments.

### Somatosensory behavior assays

Mice were acclimated to a small rectangular or round Plexiglas chamber where they could move freely but could not stand up. Selected mechanical stimuli were delivered to one hind paw when mice were calm, still, and with all four paws on the raised mesh platform. Mice were habituated to the testing chambers for one week before behavioral tests were performed. C57 and CD1 mouse lines were used, with an equal number (10) of male and female mice included. Some animals were tested multiple times, and in these cases, the quantification of behavior features was averaged across multiple trials for a given animal. Stimuli were applied to the hind paw of each mouse through the mesh floor. Cotton swab tests consisted of contact between the cotton Q-tip and the hind paw of the mouse until paw withdrawal was observed. Dynamic brush tests were performed by wiping a concealer makeup brush (e.l.f.^TM^, purchased at the CVS) across the hind paw from back to front. Light pinprick tests were performed by touching a pin (Austerlitz Insect Pins^®^) to the hind paw of the mouse. The pin was withdrawn as soon as contact was observed.

Heavy pinprick tests were performed by sharply pressing this pin into the paw so that it was pushed upwards. The pin was withdrawn as soon as approximately 1/3 of the pins length had passed through the mesh. For application of von Frey hairs (VFHs, Stoelting Company, 58011), we used 3 different forces: 0.6 grams, 1.4 grams, and 4 grams. As previously described, each VFH was directed at the center of the plantar paw and pressed upward until the filament bent^13^. For the four natural stimuli and VFHs, an animal that did not respond within 2 seconds of stimulus delivery was considered as non-responsive. For inducing chronic inflammatory pain, ~10 μL of Complete Freud’s Adjuvant, CFA (Sigma, F5881) was injected into the plantar surface of 3% isoflurane anesthetized mice as previously published^52^. For analgesic painkiller treatment, a 50 μL cocktail of meloxicam (2 mg/kg, Penn Veterinary Hospital) and buprenorphine (0.5 mg/kg, Henry Schein Animal Health, 059122) were injected subcutaneously into the back of restrained mice. Approximately 45 minutes separated injection of painkillers and behavioral testing.

### Scoring behavioral movement features

Onset of head turn is defined as a movement of the animal’s head from a stationary position towards the stimulated hind paw. Paw height and paw velocity were extracted from the high speed videos and processed with Photron FastCAM software. Paw height was scored in millimeters for the first paw withdrawal as the distance from the mesh floor to the highest point following natural or optical stimulation. Paw velocity of the first withdrawal was scored as the distance in millimeters from initial paw lift to the highest point, divided by the time in seconds between the two points. The pain score is a composite score of four individual behavior features: orbital tightening, paw shake, paw guard, and jumping. For example, if a given animal displayed one of those features, it would receive a pain score of 1. An animal was scored as displaying an orbital tightening if its eyes went from fully open to partially or fully closed following stimulus application. Paw shaking was defined as high frequency paw flinching. Jumping was defined as all four paws off the mesh floor at the same time following a stimulus application. Lastly, paw guard was defined as any abnormal placement of the paw back to the mesh floor following stimulus application.

### Peripheral optogenetics

To optically activate the nerve terminals of MRGPRD-ChR2 mice through the hind paw skin of freely behaving animals, mice were placed in the same setup and given the one-week habituation as described above for natural stimuli. All mice were scored blind to genotype and *ChR2^f/f^* littermates without the Cre-driver were used as negative controls (mice were genotyped after finishing the behavior tests). In addition, 10 mW 532 nm green laser light (Shanghai Laser and Optics Century, GL532T8-1000FC/ADR-800A) was shined to the hind paw of control and MRGPRD-ChR2 mice as another negative control. To induce a behavioral response in MRGPRD-ChR2 mice, we shined 20 mW 473 nm blue laser light (Shanghai Laser and Optics Century, BL473T8-150FC/ADR-800A) to one of the hind paws. To induce a behavioral response in TRPV1-ChR2 mice, 10 mW 473 nm blue laser light was used. For both green light control and blue light experiments we used 10 hz 20 mW sinewave light generated by a pulse generator (Agilent 10MHZ Function Waveform Generator, 33210A) connected to the laser source. For all stimulations, the laser light was delivered via an FC/PC optogenetic patch cable with a 200 μm core opening (ThorLabs, M72L01) and there was approximately 1 cm of space between the cable terminal and the hind paw skin. Light power intensity for each experiment was measured with a digital power meter with a 9.5 mm aperture (ThorLabs, PM100A). Lastly, light was only applied to mice standing on all four paws, calm and still, but not in the process of grooming.

### Tissue preparation and histology

Procedures were conducted as previously described^53^. Briefly, mice used for immunostaining were transcardially perfused with 4% PFA/PBS, and dissected tissue (either skin or spinal cord and DRGs/TGs) was post-fixed for 2 hr in 4% PFA/PBS at 4° C. Tissue used for immunostaining was cryo-protected in 30% sucrose/PBS (4% overnight) before freezing, except the c-FOS experiments where tissue was kept at room temperature and proceeded directly for vibratome sectioning. Frozen glabrous skin, DRG/spinal cord, and TG sections (20-30 mm) were cut on a Leica CM1950 cryostat. Immunostaining of sectioned TG, DRG, spinal cord, and glabrous skin tissue, was performed as described previously^53,54^. The following antibodies were used: chicken anti-GFP, 1:1000 (Aves, GFP-1020), rabbit anti-CGRP, 1:1000 (Immunostar, 24112), conjugated IB4-Alex594, 1:200 (Molecular Probes, I21411), guinea pig anti-VGLUT1, 1:1000 (Fisher, AB5905), rabbit anti-NFH, 1:1000 (Sigma, N4142), and rabbit anti-cFOS, 1:100, (Santa Cruz, sc-52).

### c-FOS staining

For c-FOS staining following optogenetic stimulation, MRGPRD-ChR2 mice were manually restrained and scuffed for 10 minutes while 10 Hz 20 mW blue light was shined directly to the ear and ear canal. We waited approximately 1.5 hours after optogenetic stimulation, and transcardially perfused the mouse with 4% PFA followed by a four hour post-fixation period. We then cut 50 μm sections with a vibratome followed by performing free-floating immunohistochemistry^53^.

### Statistical Analyses

An exploratory factor analysis with orthogonal Varimax rotation was conducted with SPSS to determine which of the initial eleven parameters contributed to at least 10% of the variance. We initially found that four parameters (total paw time, paw airtime, paw at apex, and paw time after apex) were highly correlated so we only used one (paw air-time) for subsequent analysis, leaving a total of eight parameters (Supplemental Fig. 3a). We then performed three iterations of an exploratory factor analysis using an eigenvalue cut-off of 1.0 with each being confirmed to have patterned relationships with the bartlett’s test of sphericity (p < 0.001). We then removed parameters that had either low factor loadings or cross-loaded onto multiple factors (Supplemental Fig. 3b). We considered factor loading coefficients of < 0.35 as low and not significantly contributing to a particular principle component. The first iteration revealed three Principle Components (in blue) that accounted for 62.7% of variance (Supplemental Fig. 3b – Iteration 1). Analysis of the rotated component matrix revealed that response time and head duration cross loaded onto multiple principle components so they were removed. The second iteration revealed two Principle Components (in blue) that accounted for 60.9% of variance (Supplemental Fig. 3b – Iteration 2). Analysis of the rotated component matrix revealed that full turn duration cross loaded onto multiple principle components and total time had a low factor loading (i.e., < 0.35) so they were removed. The final iteration revealed a single Principle Component (in blue) that accounted for 57.3% of variance with paw-air time having a low factor loading (Supplemental Fig. 3B – Iteration 3). We settled on three of the remaining parameters (paw height, paw velocity, and pain score).

We performed dimension-reduction with a Principle Component Analysis using the paw height, paw-air time, paw velocity, and pain score. The contributing weights, as represented by eigenvalue, for each syllable of each genotype/sex database, were determined using SAS. We could then combine normalized z-scores for each syllable into a single one-dimensional principle component for every stimulus trial. This process was conducted independently for males and females of both genotypes, generating their own set of eigenvalues for each syllable that could then be used to transform the three-dimensional data (paw height, paw velocity, and pain score) to a single dimension (Supplemental Table 1). Individual behavioral movement features, VFH filaments, and TRPV1-ChR2 and MRGPRD-ChR2 groups were compared using ANOVA followed by Tukey’s multiple comparison with p-value threshold set to 0.05.

### Machine learning

We classified paw withdrawal reflexes into “pain” and “non-pain” categories, using four measurements obtained from the high-speed imaging data: paw-air time, paw velocity, paw height and pain score. A classification pipeline consisted of the following steps (scikit-learn, 0.18.1): 1) the first principal component (PCA1) was derived from the training data, 2) the first component scores for the training data were used to train a support vector machine (SVM) with radial basis function kernels (kernel coefficient gamma = 1, penalty parameter C = 1), and 3) for a given trial, the SVM predicts the probability of the presence of a “pain” response based on that trial’s component score for the training-data PCA1. The data used to generate the PCA1 and train the SVM for each figure can be seen in Supplemental Table 2.

## Author Contribution

I.A.S, N.T.F., X.D., and W.L designed experiments. I.A.S, N.T.F., J.B., P.D., and Mark L. carried out experiments. Ming L. and L.D. performed statistics and machine learning analyses. All authors contributed to the writing and editing of the manuscript.

## Acknowledgments

We thank members of the Luo and Ma lab for helpful discussion of this work and comments on this manuscript. We thank Michael Granato for technical and intellectual consultation at the onset of this work. We also thank Dragan Vujovic, Mercedes Davis, Minah Kwon, and Mikayla Joffe for technical assistance. This work was supported by National Institutes of Health (NIH) R01 grants (NS083702 and NS094224) and the Klingenstein-Simons Fellowship Award in the Neurosciences to W.L. I.A.S. and N. T.F. are supported by NIH grant (K12GM081259), and LAS is also supported by Burroughs Wellcome Fund grant PDEP and NIH K99 grant (DE026807).

---

**Supplemental Figure 1.**
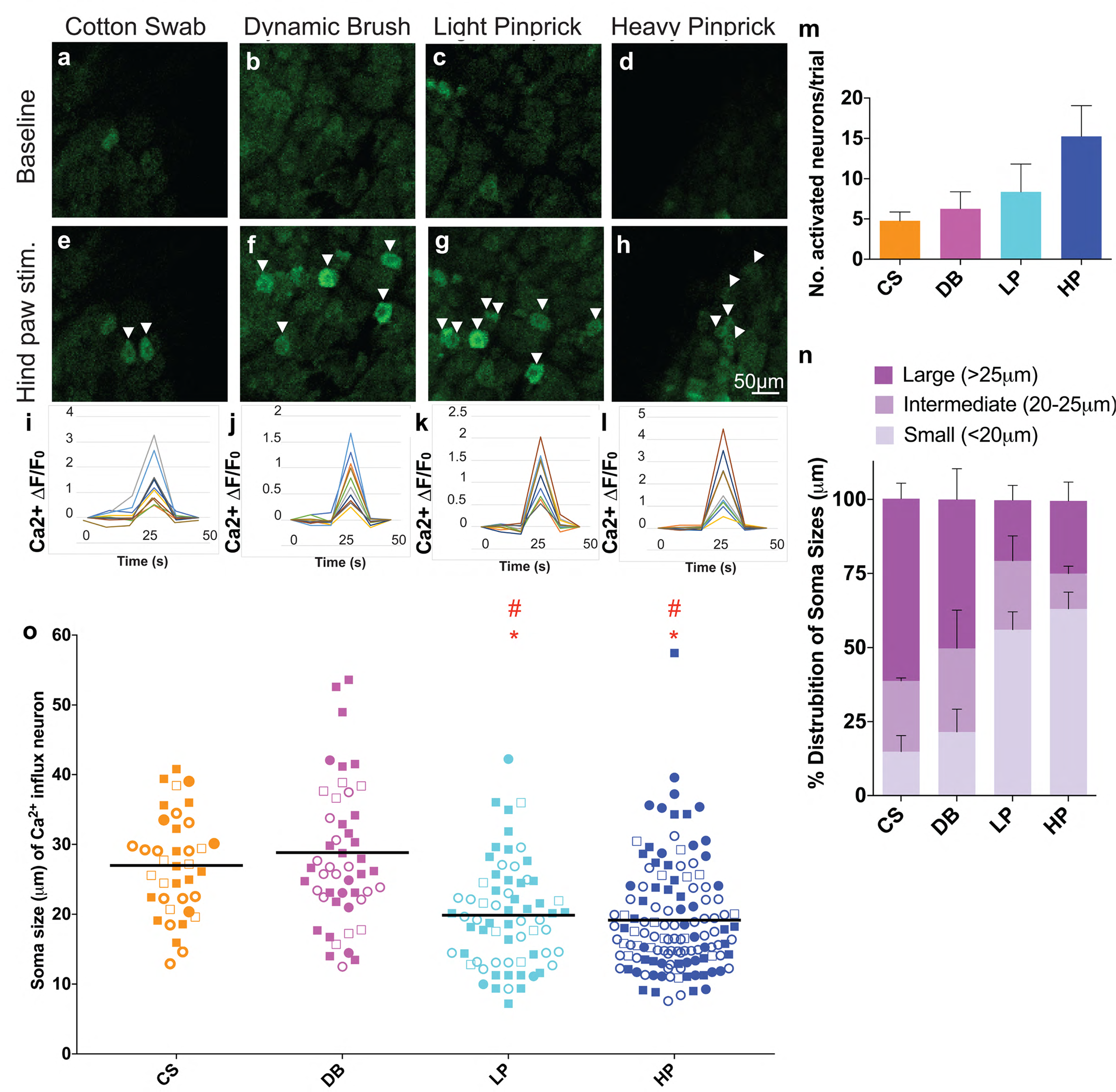
Whole animal DRG neuron calcium imaging combined with hind paw stimulation of natural mechanical stimuli. (a-h) Images of *GCAMP6* florescence before (baseline) and during hind paw stimuli of cotton swab (a, e), dynamic brush (b, f), light pinprick (c, g), and heavy pinprick (d, h). Scale bar of 50 μm the same on all images. (i-l) Example Ca^2+^ transients from 1 of 4 representative mice with each stimulus (stimulus name is directly above each graph) showing time-windows of activating neurons before, during, and after stimulus application. Note: stimulus was applied for approximately 1 second at the time-point of 24 seconds. (m) Number of activated neurons per DRG, determined by Ca^2+^ influx (ΔF/F_0_ > 20%), in 4 *Pirt-GCAMP6* mice (2 trials/mouse). (n) Soma size per activated neuron, determined by Ca^2+^ signal, in 4 *Pirt-GCAMP6* mice (2 trials/mouse). For each of 4 animals tested, the percentage of small, medium, or large diameter neurons are plotted and the 4 animals are combined here. Data are plotted according to the 4 stimuli used in panels c-j. Errors bars represent SEM. (o) Graph shows all raw data values combined for four different animals with four different stimuli - cotton swab, dynamic brush, light pinprick, and heavy pinprick. The four different animals are distinguished by either open circles, closed circles, open squares, or closed squares. Each shape represents a single neuron. Red asterisks represent p-values <0.05 when comparing CS to LP or CS to HP (LP or HP > CS), while red stars represent p-values <0.05 when comparing DB to LP or DB to HP (LP or HP > CS).

**Supplemental Figure 2.**
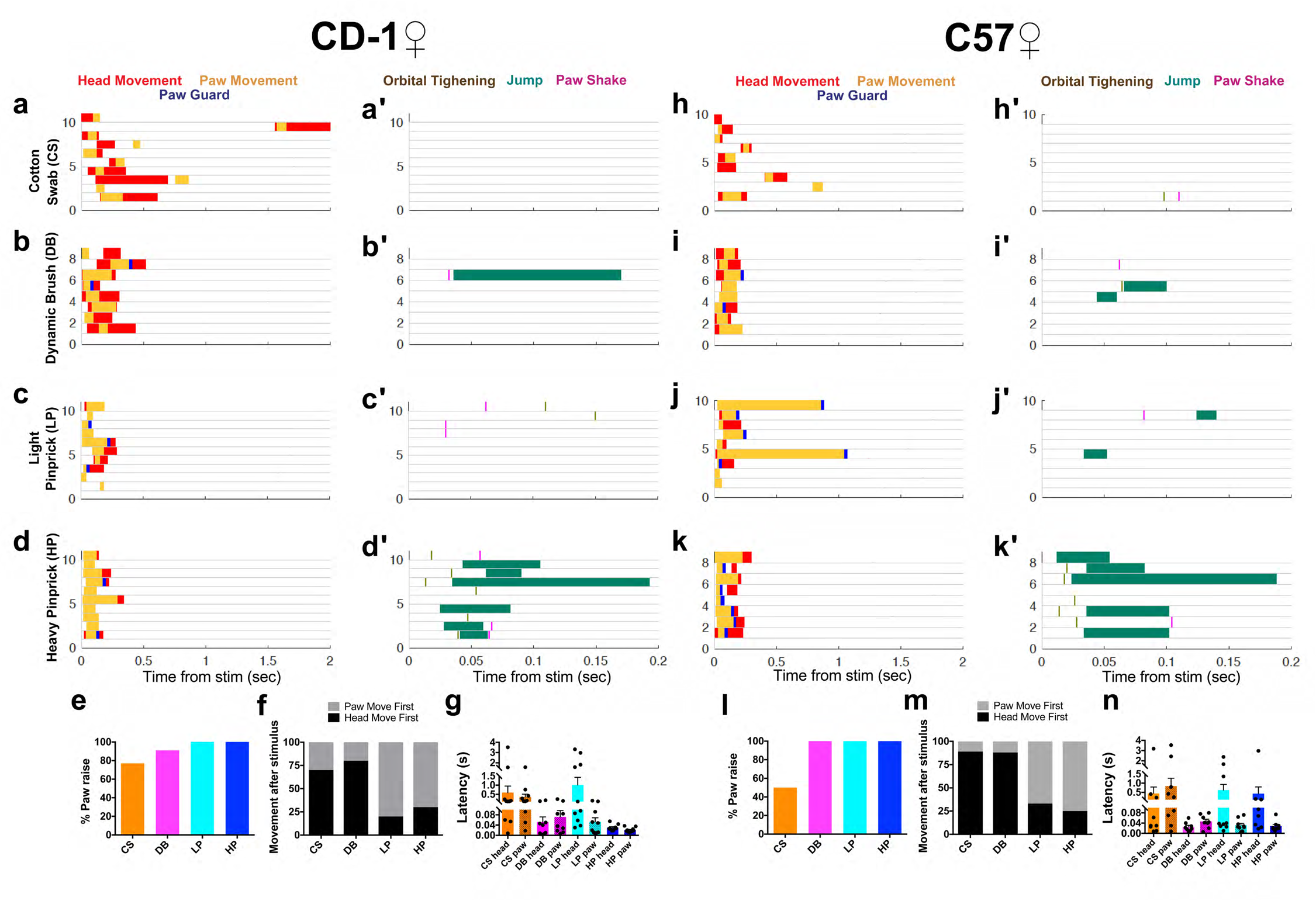
Sub-second temporal mapping of female mouse behavioral features in response to paw application of natural mechanical stimuli. Responses of CD1 and C57 female mice to paw stimulation of cotton swab (CS), dynamic brush (DB), light pinprick (LP), and heavy pinprick (HP) are plotted as raster plots, showing when six behavior features (color-coded in the figure) occurred after stimulus onset within the first 2 s (a-d, h-k) or the first 200 ms (a′-d′, h′-k′). For each raster plot, the times when the behaviors occurred are shown on the X-axis, while the Y-axis and each horizontal line show a single trial/animal. Each horizontal line from the two columns of raster plots for a given strain is from the same trial. (e) Percentage of paw raise towards a given stimuli for CD1 females, n = 10. (f) First movement, whether head (black) or paw (grey), after stimulus application for CD1 females. (g) Latency of head and paw movement upon each stimulation for CD1 females. (l) Percentage of paw raise towards a given stimuli for C57 females, n = 10. (m) First movement, whether head (black) or paw (grey), after stimulus application for C57 females. (n) Latency of head and paw movement upon each stimulation for C57 females.

**Supplemental Figure 3.**
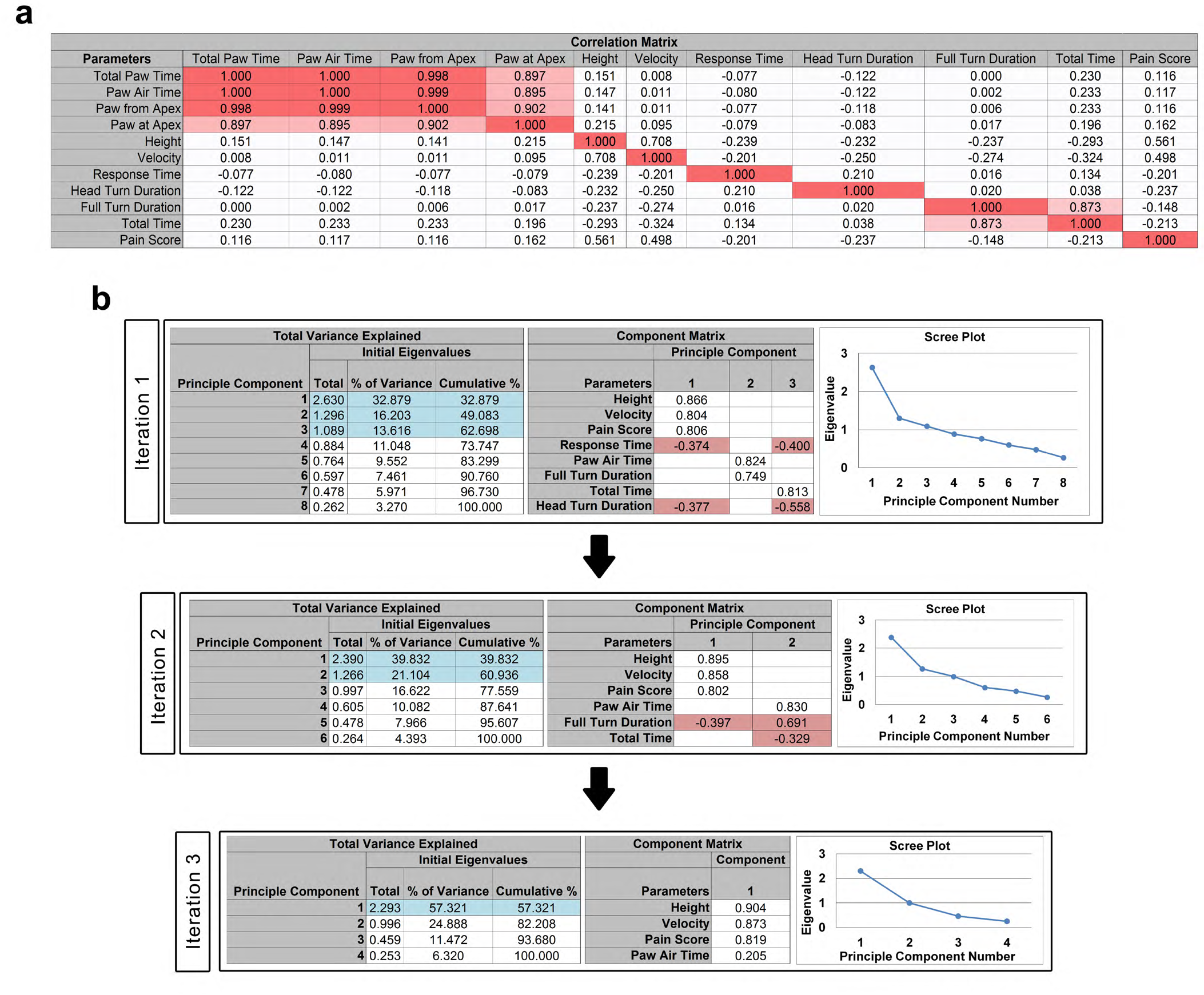
Exploratory Factor Analysis reveals four parameters that account for majority of variance. Data is derived from ~50% of trials from both C57 and CD1 males. (a) Correlation matrix between all 11 movement parameters. Data-cells marked by a gradient of red indicate correlations above 0.75. (b) Iterative exploratory factor analysis. Each iteration has three panels: 1) Cumulative “Total Variance Explained” for each principle component. Principle components with eigenvalue greater than 1.0 are highlighted blue. 2) “Component Matrix” with factor loadings for each parameter that makes up the associated principle component highlighted blue. 3) Scree plot for each principle component. Note that each iteration removes parameters that either have a low factor loading (< 0.35) or cross-load onto multiple factors (highlighted red).

**Supplemental Figure 4.**
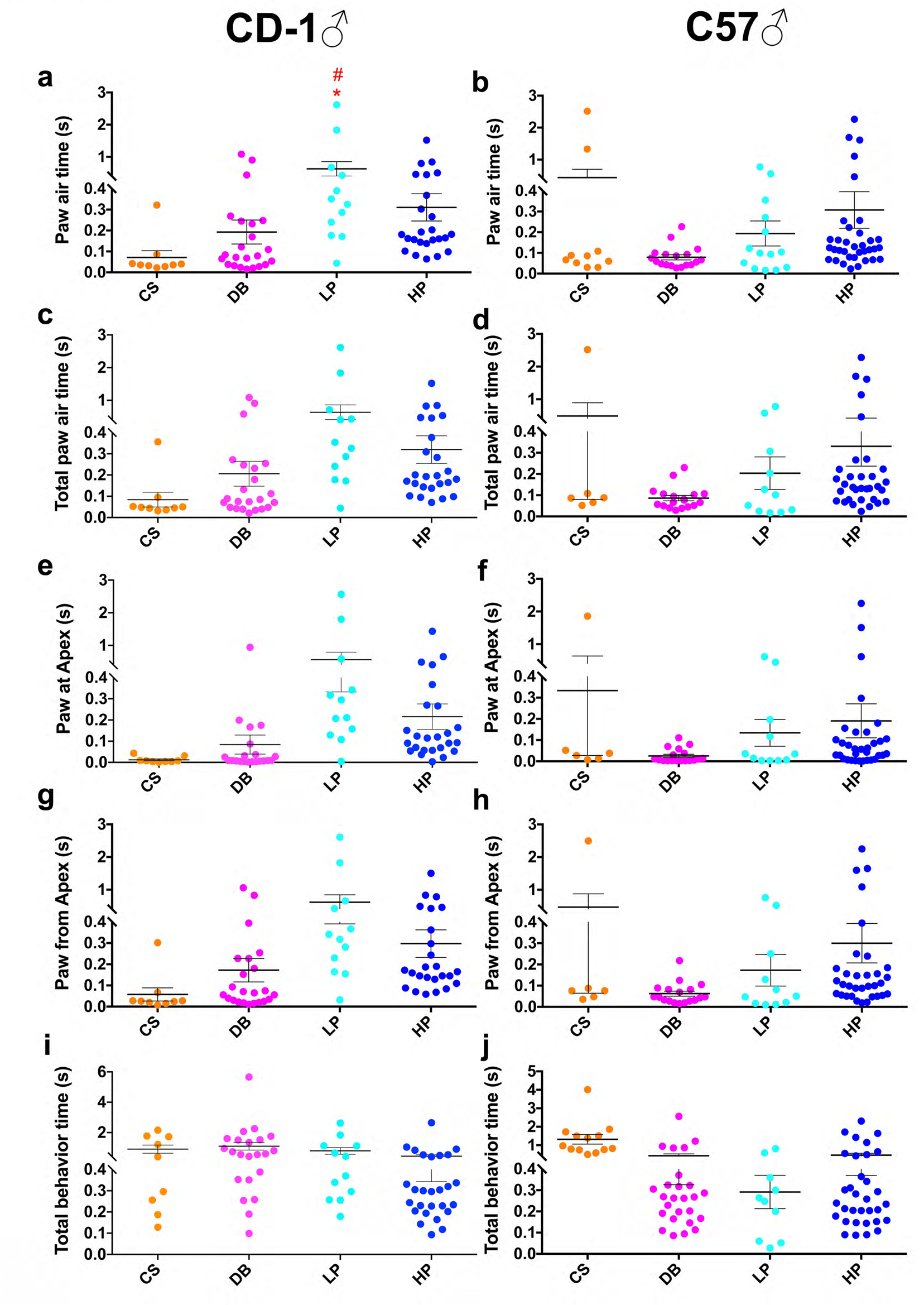
Stimulus-evoked movement features excluded from further analyses with PCA and SVM. (a, b) Paw air time measurements for CD1 males (a) and C57 males (b) after application of the four natural stimuli mentioned in the main text. Paw air time refers to the time when the animal’s stimulated hind paw is in the air. (c, d) Total paw air time measurements for CD1 males (c) and C57 males (d) after application of the four natural stimuli mentioned in the main text. Total paw air time refers to the time when the animal’s stimulated hind paw is in the air, including the time when it first moves in paw before lifting away from the surface. (e, f) Paw at apex measurements for CD1 males (e) and C57 males (f) after application of the four natural stimuli mentioned in the main text. Paw at apex refers to the time when the animal’s stimulated hind paw is held at its maximal point in the air. (g, h) Paw from apex measurements for CD1 males (g) and C57 males (h) after application of the four natural stimuli mentioned in the main text. Paw from apex refers to the time when the animal’s stimulated hind paw is coming down from its maximal height towards placement back on the surface. (i, j) Total behavior time measurements for CD1 males (i) and C57 males (j) after application of the four natural stimuli mentioned in the main text. Total behavior time refers to the time the animal first begins a movement (either head turn or paw lift) until the time these movements are completed.

**Supplemental Figure 5.**
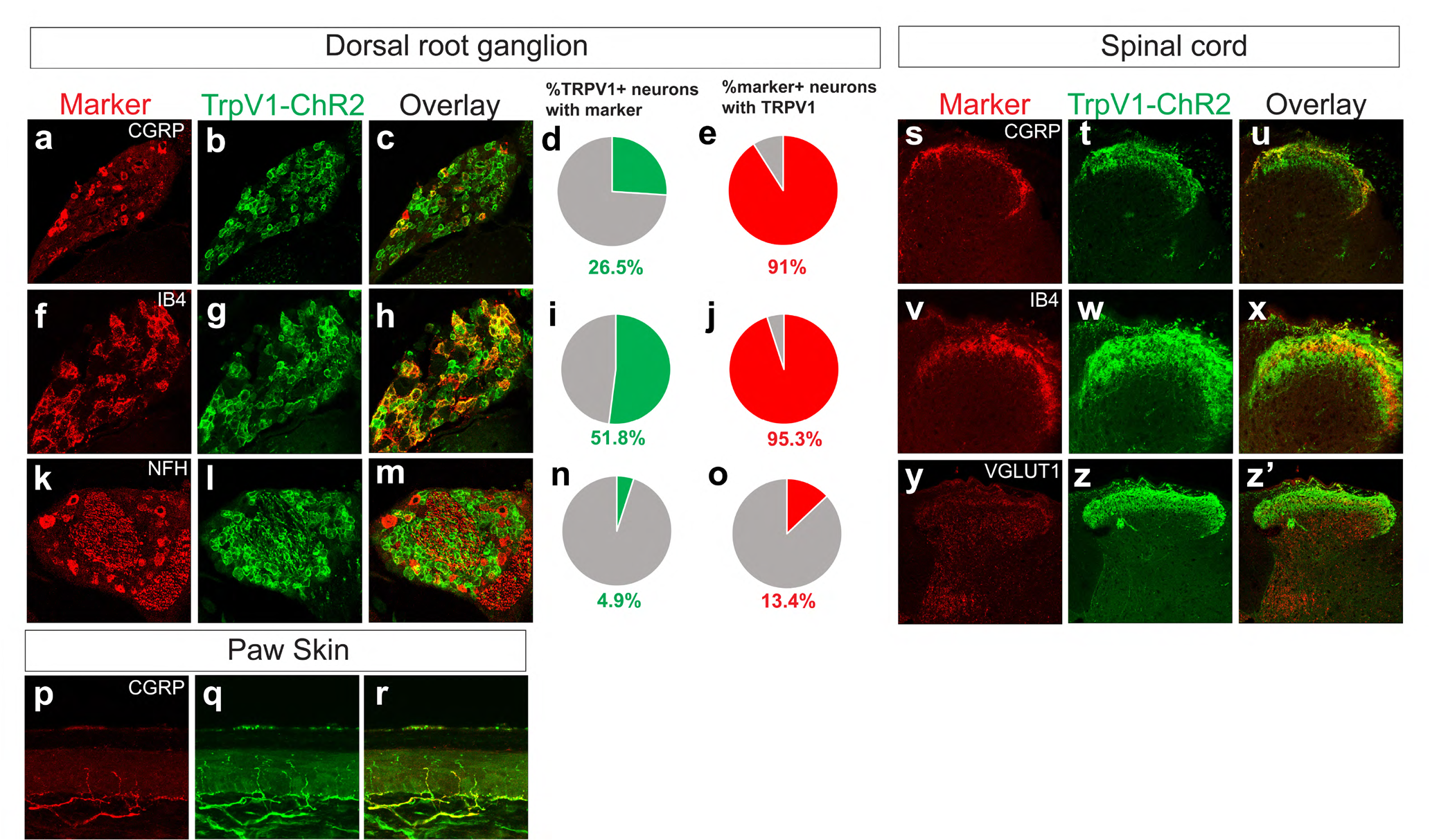
Histology of the dorsal root ganglia, spinal cord, and plantar paw skin of *TrpV1^Cre^; Ai32* mice. (a-c) DRG sections immunostained with antisera directed against GFP, which recognize ChR2-EFYP fusion protein, and CGRP, and the percentages of overlap are shown with bar graphs (d, e). Corresponding double immunostaining was done on spinal cord tissue (s-u). (f-h) DRG sections immunostained with IB4 and antiserum directed against GFP, and percentages of overlap are shown with bar graphs (i, j). Corresponding double immunostaining was done on spinal cord tissue (v-x). (k-m) DRG sections immunostained with antisera directed against GFP and NFH and percentages of overlap are shown with bar graphs (n, o). Corresponding double immunostaining was done on spinal cord tissue with antisera against GFP and VGLUT1 (y-z’). Plantar paw skin double immunostaining was performed with antisera directed against GFP and NFH (p-r). n = 3 mice between P21-P28 for histology with DRG, spinal cord, and plantar paw skin.

**Supplementary Figure 6.**
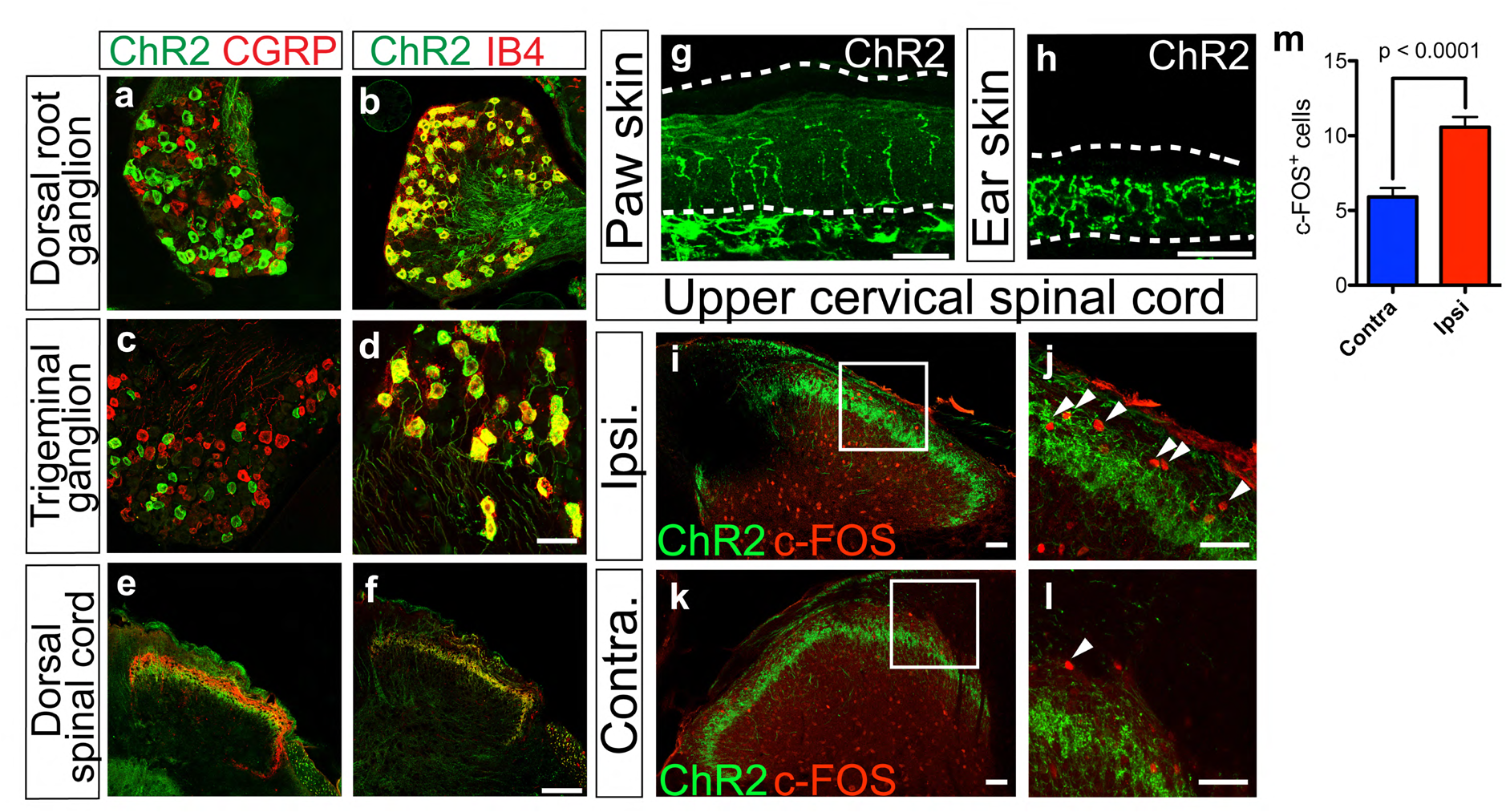
Genetic targeting and activation of ChR2 in MRGPRD^+^ neurons. (a, b) Representative confocal image of immunostaining with dorsal root ganglia sections of *MrgD-ChR2* mice, showing no overlap between DRG neurons expressing ChR2-EYFP and CGRP, but complete overlap between those expressing ChR2-EYFP and binding IB4. (c, d) Similar expression patterns are observed after immunostaining with the trigeminal ganglia sections for antisera that detect ChR2-EYFP, CGRP, and IB4. (e, f) Immunostaining with the dorsal spinal cord sections of *MrgD-ChR2* mice showing efficient targeting of ChR2-EYFP to central terminals that do not overlap with CGRP+ (e) but IB4+ (f) central terminals. (g, h) Immunostaining showing efficient targeting of ChR2-EYFP to peripheral terminals in the dermal plantar paw (g) and ear skin (h) of *MrgD-ChR2* mice. (i-l) Immunostaining of c-FOS with the upper cervical spinal cord sections following optogenetic ear stimulation of *MrgD-ChR2* mice shows increased number of c-FOS^+^ neurons in the ipsilateral (blue light) superficial dorsal horn (i, j) compared to the contralateral side (no light) (k, l). j and l are magnified from the white box areas in I, k. (m) Quantification of c-FOS cells following optogenetic stimulation of MrgD-ChR2 mice. P-value is from student’s t-test and error bars represent SEM. n = 3 mice. Scale bars are 50 μm.

**Supplemental Figure 7.**
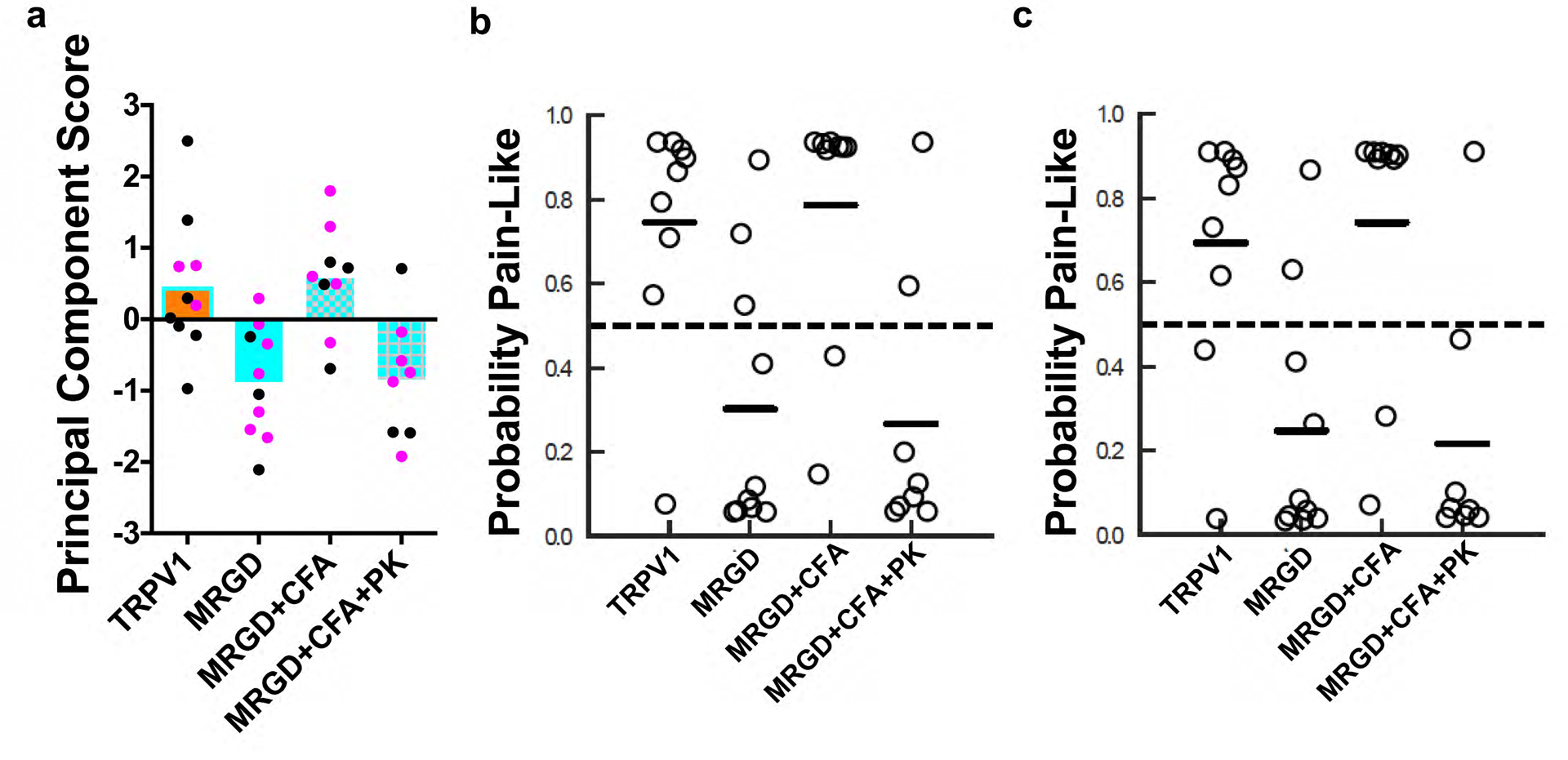
First Principal Component (PC) scores and SVM predictions for optogenetic paw withdrawals in a sex specific manner. (a-c) Each dot represents a single animal. (a) PC scores of TRPV1-ChR2 and MRGPRD-ChR2 mice at baseline, after CFA, and CFA + painkillers. Eigenvectors derived from wild type male mice (C57+CD1) and represented by black dots, or eigenvectors derived from wild type female mice (C57+CD1) and represented by magenta dots. (b, c) SVM pain-probability graphs using wild type male mice (C57+CD1) (b) or wild type female mice (C57+CD1) (c) as training datasets, to predict the probability of a pain response for TRPV1-ChR2 and MRGPRD-ChR2 optogenetic responses in baseline, after CFA, and CFA + painkillers.

**Supplemental Figure 8.**
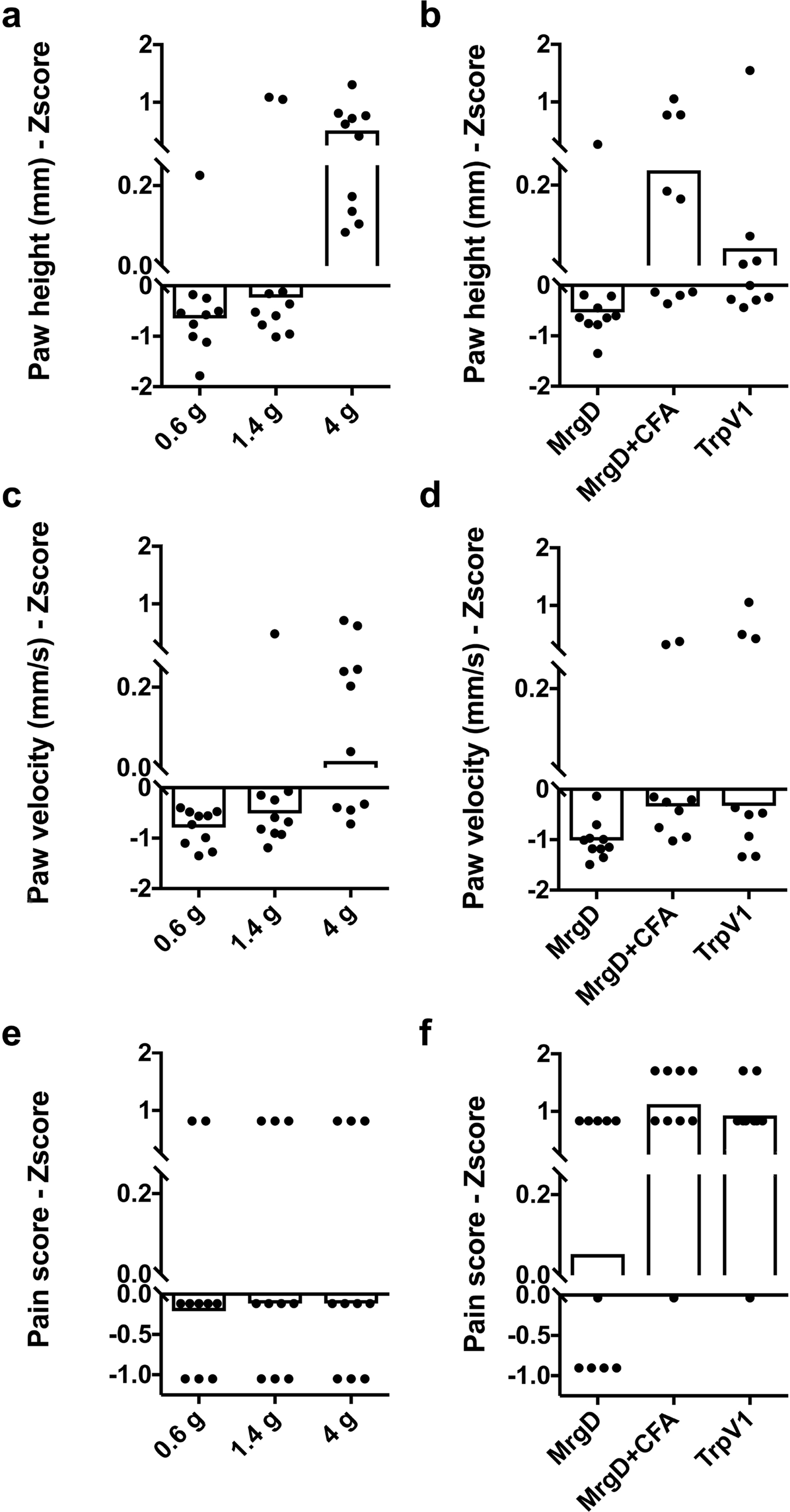
Z-scores for paw height and velocity, and pain score comparing Von Frey hairs with optical stimuli. Data are compiled and plotted the same as Figure 3. Shown are the Z-score measurements for paw height (a, b), paw velocity (c, d), and pain score (e, f). (a, c, e) are measurements from Von Frey hairs at 0.6 g, 1.4 g, and 4 g and correspond to the data shown in Figure 5. (b, d, f) are measurements from TRPV1-ChR2 mice (baseline) and MRGD-CHR2 mice (baseline and post-CFA) and correspond to the data shown in Figure 6.

**Supplemental Table 1.** Eigenvalues for calculating the first Principal Component (PC) scores. These values were determined using SAS software using the Supplementary Raw Data File 2.

| Variable | C57 Male | C57 Female | CD1 Male | CD1 Female | C57+CD1 Females | C57+CD1 Males | Both sexes + strains |
| --- | --- | --- | --- | --- | --- | --- | --- |
| Z Paw Velocity | 0.57742 | 0.584451 | 0.57431 | 0.594993 | 0.590982 | 0.568939 | 0.582674 |
| Z Paw Height | 0.605828 | 0.560678 | 0.591051 | 0.570744 | 0.564709 | 0.607105 | 0.580398 |
| Z Pain Score | 0.547319 | 0.586564 | 0.566416 | 0.565892 | 0.576059 | 0.554736 | 0.568885 |

**Supplemental Table 2.** Data used to generate and train the SVM to make predictions about pain-like probabilities.

Data used for support-vector machine learning
| PCA Datasets | Fitting Datasets | Figure |
| --- | --- | --- |
| CD1(M) | CD1(M) - CS,HP | 4 A-D |
| CD1(F) | CD1(F) - CS,HP | 4 E-H |
| C57(M) | C57(M) - CS,HP | 4 I-L |
| C57(F) | C57(F) - CS,HP | 4 M-P |
| CD1(M) | CD1(M) - CS,HP | 5 G |
| CD1(M), CD1(F), C57(M), C57(F) | CD1(M), CD1(F), C57(M), C57(F) - CS, HP | 6 I |
| CD1(M), CD1(F), C57(M), C57(F) | CD1(M), C57(M) - CS, HP | Supp. 7B |
| CD1(M), CD1(F), C57(M), C57(F) | CD1(F), C57(F) - CS, HP | Supp. 7C |

**Supplemental Raw Data File 1.Ca^2+^ transients (ΔF/F_0_) of activated DRG neurons with whole animal imaging**. These arbitrary ImageJ values show the time-windows before, during, and after hind paw stimulation of cotton swab, dynamic brush, light pinprick, and heavy pinprick. Time-point 4, which corresponds to ~24 seconds into imaging, is when the stimulus was applied. Mean refers to an individual neuron.

**Supplemental Raw Data File 2. Behavioral Parameter Raw Data**. Paw air time, paw velocity, paw height, and pain score for each trial for CD1 male, CD1 female, C57 male, C57 female, MrgD-ChR2, and Trpv1-ChR2 mice as well as VFH for CD1 male mice. Z-scores and PCA1s are also plotted.

